# *BrainGENIE*: The Brain Gene Expression and Network Imputation Engine

**DOI:** 10.1101/2020.10.27.356766

**Authors:** Jonathan L. Hess, Thomas P. Quinn, Chunling Zhang, Gentry C. Hearn, Samuel Chen, Neuropsychiatric Consortium for Analysis and Sharing of Transcriptomes, Sek Won Kong, Murray J. Cairns, Ming T. Tsuang, Stephen V. Faraone, Stephen J. Glatt

## Abstract

*In vivo* experimental analysis of human brain tissue poses substantial challenges and ethical concerns. We developed a novel method called the Brain Gene Expression and Network Imputation Engine (*BrainGENIE*) that uses peripheral-blood transcriptomes to predict brain-tissue-specific gene-expression levels. *BrainGENIE* reliably predicted brain-tissue-specific expression levels for 1,733 – 11,569 genes (false-discovery rate-adjusted *p*<0.05), including many transcripts that cannot be predicted reliably by a transcriptome imputation method such as *PrediXcan*. We tested the generalizability of *BrainGENIE* in external within-individual data from *ex vivo* peripheral blood and *postmortem* brain samples from the Religious Orders Study and Memory and Aging Project, wherein we validated 39% of predicted gene expression levels as concordant with observed expression levels in dorsolateral prefrontal cortex and 23% in caudate. *BrainGENIE* recapitulated diagnosis-related gene expression changes in brain better than direct correlations from blood and predictions from *PrediXcan. BrainGENIE* complements and, in some ways, outperforms existing transcriptome-imputation tools, providing biologically meaningful predictions and opening new research avenues.

## Introduction

Brain disorders cause considerable disability worldwide ^1^. Typically, *in vivo* molecular assessment of human disease centers on the primarily affected tissue(s) or the site of pathogenesis, but that is not possible for brain disorders unless neurosurgical intervention is required. Collecting *ex vivo* human brain tissue in an experimental setting for neuropsychiatric research is infeasible given the considerable risks associated with brain biopsy. There are numerous research questions that would be answered best by studying living human brain tissue, but which therefore remain unaddressed. Transcriptome imputation offers a non-invasive alternative to brain biopsy by allowing investigators to infer tissue-specific gene expression without directly assaying gene-expression levels. Instead, brain-tissue-specific predictions about gene expression can be made in living persons by proxy of genetic variants genotyped from peripheral tissues (*e*.*g*., blood), capitalizing on the known regulatory relationships between those variants and the expression of transcripts.

*FUSION* and *PrediXcan* are two software tools that model tissue-specific effects of expression quantitative trait loci (eQTLs) on the expression of proximal genes (*cis*-eQTLs) in order to impute transcriptome profiles. These methods have been successful in prioritizing genome-wide association study (GWAS) hits and have helped reveal putative mechanisms underlying complex disorders ^2–6^. With both methods, there is a striking disparity between the number of genes imputable in the brain *versus* tissues outside of the central nervous system (CNS): to wit, *FUSION* imputes an average of 3,158 genes in brain (range=1,604 – 5,855 across 12 brain tissues (including a pair of re-sampled tissues from frontal cortex [BA9] and cerebellum) compared with 5,592 in non-CNS tissues; similarly, *PrediXcan* imputes an average of 4,337 genes in the brain (range=2,559 – 6,794) compared with 6,262 genes (range=1,642 – 10,012) outside the CNS. Furthermore, the majority of genes in the brain transcriptome are not reliably predicted by either *FUSION* or *PrediXcan*, suggesting that a large amount of variance in transcriptome profiles cannot be captured by eQTLs alone. A recent addition to the suite of genotype-based transcriptome-imputation methods is *TIGAR*, which uses a Bayesian modeling framework for predicting gene expression from eQTL data. Using a data-driven nonparametric model of *cis*-eQTL signals, *TIGAR* further increases the number of imputable genes, and improves accuracy of predictions ^7^. A Bayesian hierarchical model called *EpiXcan* builds upon *PrediXcan* by applying epigenetic annotations to optimize the weights assigned to *cis*-eQTLs and increase predictability of gene-expression levels ^8^.

Tissue-specific and tissue-dependent gene expression helps differentiate between brain and peripheral tissues, but compelling evidence also shows that brain and blood exhibit comparable transcriptome profiles ^9–12^. Our group systematically reviewed relevant literature on this topic, and found that gene expression profiles in blood and brain are moderately correlated (Pearson’s *r* of 0.24 – 0.64), with 35 – 80% of genes expressed in both tissues ^9^. In a later study, we found empirical evidence that ∼90% of weighted gene-gene interaction networks identified in peripheral blood transcriptomes were significantly preserved in the prefrontal cortex ^10^. Brain and blood also show significant overlap with respect to eQTLs ^11,12^, signifying that shared genetic effects (albeit with small effect sizes) may in part explain the comparability of gene expression in blood and brain. Data from previous studies suggest that blood gene expression is a reasonable (albeit imperfect) direct proxy for brain gene expression. Another advantage of capitalizing on human blood transcriptomes for brain gene-expression imputation is that such data are widely available in public repositories and also can be generated *de novo* with relative ease and cost-effectiveness. Unlike DNA variants, transcriptome profiles in blood fluctuate over time, and they may reflect valuable information about corresponding temporal changes in the brain throughout development or over the course of an exposure or intervention. Characterizing temporal changes in tissue-specific gene expression is not possible with existing *cis*-eQTL-based toolsets.

Based on this evidence and logic, we sought to capitalize on the transcriptome similarity between brain and blood (and the easy accessibility of blood) to make predictions about gene expression in the brain solely based on observed expression in the periphery. Simultaneously, we sought to develop an expression-based transcriptome-imputation method that complements existing *cis*-eQTL-based transcriptome-imputation methods. We achieved these goals with a novel Brain Gene-Expression and Network-Imputation Engine (*BrainGENIE*), which imputes brain-tissue-specific gene-expression profiles based on gene-expression profiles assayed from peripheral blood. *BrainGENIE* is implemented in the *R* statistical environment and is distributed as freely available software (https://github.com/hessJ/BrainGENIE).

*BrainGENIE* is not the first or only cross-tissue transcriptome-imputation method, but it has unique strengths that compare quite favorably with other approaches. Tissue Expression Estimation using Blood Transcriptome (*TEEBoT*), like *BrainGENIE*, uses principal components of peripheral blood transcriptomes to predict transcriptomes of other tissues ^13^. *TEEBoT* was developed using an earlier release of GTEx (v.6) data; hence, its modeling of brain-tissue-specific transcriptomes was limited to cerebellum and caudate due to sample-size restrictions. In contrast, *BrainGENIE*, which was built on GTEx (v.8), imputes gene-expression levels in 12 different brain tissues. *B-GEX* similarly achieved a limited modeling of brain regional transcriptomes due to sample-size restrictions as it uses an older version of GTEx that included fewer donors with paired blood and brain data. Moreover, *B-GEX* models use individual gene transcripts from blood selected using a Bayesian regression to predict brain gene expression, capturing less variance than principal components, and thus limiting predictive power. In short, because *BrainGENIE* was built on a better dataset, models more brain tissues, and uses principal components of whole-blood gene expression, it matches the strengths of competing methods and overcomes some of their limitations. Since there is no equivalent blood-based transcriptome-imputation method available that has modeled all regions of the brain like *BrainGENIE*, we benchmarked *BrainGENIE* against *PrediXcan*. The two methods are conceptually similar and there are data available from *PrediXcan* for all 12 brain tissues modeled by *BrainGENIE*, enabling direct comparisons. As such, we used the methodology of *PrediXcan* as a basis for developing *BrainGENIE* so that the results from the two methods could be directly compared. Comparing these two methods helped us to understand differences (and points of convergence) between the use of blood-based gene-expression profiles *versus* eQTLs to impute region-specific gene expression in the brain, and illuminated the relative strengths and weaknesses of each approach.

## Methods

### Data

RNA-sequencing (RNAseq) data were obtained from *postmortem* brain tissue and whole blood of 267 adult human donors. These data were generated as part of the GTEx Project (v.8), and were downloaded from the dbGaP repository (phs000424.v8.p2). Tissues selected for the GTEx Project were from donors free of brain pathology. The inclusion criteria for donors were as follows: a body mass index (BMI) of 18.5 – 35, tissue collected within 24 hours of time of death, no blood transfusions within 48 hours of time of death, no metastatic cancer, no chemotherapy or radiation within two years of time of death, and no communicable diseases that would disqualify donors from tissue donation. Transcriptome profiles were available for 12 brain tissues, two of which were collected at the same time as the donor non-brain tissue and preserved in PAXgene tissue kits (cerebellum and frontal cortex), and 10 tissues were collected and flash frozen at the University of Miami Endowment Brain Bank from whole brains that were left unfixed and shipped on wet ice (amygdala, anterior cingulate cortex, caudate, cerebellum [re-sampling], frontal cortex [re-sampling], hippocampus, hypothalamus, nucleus accumbens, putamen, and substantia nigra). Genotype-based principal components that captured genetic ancestry were available for 218 donors. Some brain tissues were not available from some donors; hence, pairs of blood-brain transcriptome profiles ranged from *n*=86 – 153. Ages of donors ranged from 20 – 70 years old with the distribution skewed toward older persons (50% over the age of 61 years, mean age=57.9 years). Approximately 30% of donors were female. Donors were recorded predominantly as of European-American ancestry (*n*=199, 91%), with 8% recorded as African American (*n*=17), and less than 1% recorded as Asian, Alaska Native, or American Indian (*n*=2).

### Normalization of RNAseq Data from GTEx

Total RNA was extracted from whole blood stored in PAXgene Blood RNA (Qiagen®) tubes. Total RNA was extracted from PAXgene fixed tissues sampled from frontal cortex and cerebellum at the same time as the donor non-brain tissues. Whole brains had been shipped on wet ice to the University of Miami Endowment Brain Bank where 10 brain tissues were sampled (including re-sampled frontal cortex and cerebellum as close to original site) and flash frozen. All RNA samples that were used for RNAseq had RNA integrity number (RIN) values ≥ 6.0 as measured by Agilent Bioanalyzer. Additional details related to tissue collection, library preparation, and sequencing were previously described by the GTEx Consortium ^14^. Gene-level read counts were summarized based on the Gencode 26 (GRCh38) transcript model. We took the following steps pre-process gene-counts for analysis, including: retain genes with > 0.1 read per kilobase per million (RPKM) and ≥5 read counts in at least 10 donors, quantile-normalize RPKM to adjust for between-sample variation (*limma* v.3.24.3) ^15^, and inverse-rank normalization.

### Statistical Deconvolution of Whole Blood Transcriptomes

*CIBERSORT* (v.1.04) was used to estimate the relative abundance of circulating blood-cell subtypes across GTEx donors based on expression of cell-type-specific marker genes ^16^. *CIBERSORT* uses a support vector regression model to predict abundance of cell types based on prior knowledge of cell abundance and gene-expression levels in a reference panel. For this analysis, we used a reference panel referred to as “LM22”, which was comprised of expression levels for 547 genes and measured abundances of 22 human hematopoietic cells. We then performed principal components analysis (PCA) on the profiles generated by *CIBERSORT* to identify a smaller number of components that explain variation in leukocyte abundances across GTEx donors.

### Adjusting for Confounding Variation

Covariates used by the GTEx Project in their eQTL analyses (including genotyping principal components, sequencing platform, sequencing protocol, and inferred hidden factors of gene expression profiles) were downloaded from the GTEx Project website ^17,18^. A methodological difference between *BrainGENIE* and *PrediXcan* is that we did not include “probability estimates of expression residuals” (PEER) as covariates in our multiple-regression models in order to preserve variation in transcriptome levels for model training. The GTEx Consortium cautioned that the use of PEER factors may be overly aggressive, citing that PEER factors have a large impact on variation in gene expression levels (58 – 78%) ^18^, which may remove useful biological variation. In light of this concern, we used clearly defined technical and biological covariates instead of PEER factors when modeling transcriptome profiles. Multiple regression was used to compute *residual expression values* for blood and brain tissues by removing additive effects attributed to age, sex, ancestry covariates attributed to the top three genotype-derived principal components, PCR method used in library preparation, RNAseq platform, RIN, *postmortem* interval, and death classification of each donor based on the 4-point Hardy scale ^19^. For whole blood, we included three additional covariates (defined as the top 3 PCs from *CIBERSORT*) to adjust for between-donor variation in circulating leukocyte abundance (**Supplementary Figure 1**). The residualized data, having had confounding variation partialled out, were used for building prediction models of brain gene expression.

### Training and Evaluation of BrainGENIE

We performed a single 10-fold cross-validation to estimate the predictive performance of *BrainGENIE* separately for each brain region. Paired blood-brain transcriptome profiles from GTEx donors were randomly assigned to the 10 folds. For each *training subset*, a PCA was performed on normalized blood transcriptome profiles, and linear regression was trained to predict brain-tissue-specific expression levels *per-gene* using the top *k=*5 (11% variance explained), *k=*10 (41% variance explained), *k=*20 (58% variance explained), and *k=*40 PCs (80% variance explained) of whole-blood gene expression (resulting in fold={1…10} by *k*={5,10,20,40} by gene={1…*n*_genes_} linear models). Our initial work uncovered that prediction accuracies achieved by linear regression were as good as or better than elastic net regression (the model used by *PrediXcan*); linear regression is also computationally faster to train, thus was the chosen model for *BrainGENIE*. The trained models were then deployed in the validation set to estimate the predictive performance on unseen data. The metric for prediction performance was the coefficient of determination for observed and predicted *per-gene* expression levels (*R*^2^) in the hold-out fold. This process was repeated until each fold was used as the validation set, and *per-gene* prediction performance was averaged over the validation sets. In order to have a reasonable side-by-side comparison between *BrainGENIE* and *PrediXcan*, we adopted the same criterion for “reliably predicted” as adopted by *PrediXcan*; *i*.*e*., genes that could be predicted with a cross-validation *R*^2^≥0.01 and with Benjamini-Hochberg false-discovery rate adjusted *p*-value (FDR)<0.05. When comparing all models, the 20-PC *BrainGENIE* model exhibited the best performance in the training data in terms of average *R*^2^ and number of genes with reliably predicted gene expression levels, and was selected as the final model to deploy on the external test set (described below).

### Accuracy of BrainGENIE versus PrediXcan

We used two-tailed *t*-tests (alpha = 0.05) to compare the prediction accuracy of *BrainGENIE* and *PrediXcan* as indexed by the Pearson’s *r* coefficients for genes that met the criterion for being “reliably predicted”. Tests were performed separately for each of the 12 brain tissues, and Benjamini-Hochberg FDR corrections were applied to resultant *p*-values to adjust for multiple testing. In addition, we used Pearson’s correlation tests to assess the similarity of prediction accuracies between *BrainGENIE* and *PrediXcan* for genes that both methods reliably predicted.

### Enrichment of Synaptic Gene-Sets

The aim of this analysis was to test for a difference between *BrainGENIE* and *PrediXcan* in the level of enrichment of synaptic ontologies among genes reliably predicted by these methods. Synaptic ontologies were chosen to provide another means of comparing methods, and to limit the scope of analysis to gene-sets that are fundamental to the brain, many of which are overrepresented with common genetic variants associated with neuropsychiatric disorders ^20^. Annotations for synaptic genes were obtained from the SynGO database (release 2018-07-31) ^20^. We used a two-proportions *χ*^2^ test to compare *BrainGENIE* and *PrediXcan* on the proportions of presynaptic genes (*n* genes=747) and postsynaptic genes (*n* genes=1,351) reliably predicted by each method.

### Enrichment of Brain Cell-Type Marker-Genes

The aim of this analysis was to test for a difference between *BrainGENIE* and *PrediXcan* in the level of enrichment of brain cell-type marker-genes that were reliably predicted by these methods. We obtained lists of cell-type marker-genes for neurons, astrocytes, endothelial cells, oligodendrocytes, microglia, and oligodendrocyte precursor cells identified in *postmortem* human brain samples from the Allen Brain Atlas using the *R* package *BRETIGEA* (v1.0.0) ^21,22^. Two-proportions *χ*^2^ test were used to compare *BrainGENIE* and *PrediXcan* on the proportions of marker-genes for the six brain-cell types reliably predicted by each method.

### Enrichment of Cross-Disorder Pleotropic Gene-Sets

The goal of this analysis was to employ gene-set enrichment to determine if *BrainGENIE* and *PrediXcan* differed in their ability to reliably predict gene-sets that show significant association with major neuropsychiatric disorders by GWAS. Gene Ontology (GO) identifiers were obtained for 45 gene-sets identified by GWAS meta-analysis as having a shared association across eight neuropsychiatric disorders ^23^. GO identifiers were annotated with HGNC gene symbols using the Molecular Signatures Database (v.6.2) ^24^. We calculated the proportion of genes per gene-set that were reliably predicted using *BrainGENIE* and compared this with the proportion of each gene-set reliably predicted using *PrediXcan* using a two-proportions *χ*^2^ test.

### Concordance with Disease-Related Transcriptomic Signatures

To determine how well *BrainGENIE* captures brain-relevant signatures for disease, we performed a Pearson’s correlation analysis to determine the concordance of imputed differential-expression signatures for schizophrenia (SCZ), bipolar disorder (BD), major depression (MDD), and autism spectrum disorder (ASD) derived by *BrainGENIE* with “ground-truth” differential expression signatures from *postmortem* brain published by the PsychENCODE Consortium ^25^. For this analysis, we deployed *BrainGENIE* models on completely independent blood-based transcriptome datasets for SCZ ^26–31^ (*k* studies=7, *n* cases=258, *n* controls=241), BD ^31–36^ (*k* studies=6, n cases=84, *n* controls=101), and ASD ^37–44^ (*k* studies=5, *n* cases=584, *n* controls=431). Descriptions for each dataset are provided in **Supplementary Table 1**. Pre-processing and normalization steps used to prepare blood-based transcriptome profiles for the SCZ, BD, and ASD datasets are described in our previously published studies ^10,45,46^. Details of our normalization procedure are available in the supplement (**Supplementary Methods**). The combined set of peripheral blood transcriptome data for each disorder was then supplied to *BrainGENIE* in order to impute transcriptome profiles for the frontal cortex using the 5-, 10-, 20-, and 40-PC models.

We calculated differential gene expression (DGE) in blood between affected cases and unaffected comparison individuals *via* combined-samples mega-analyses, adjusting for covariates commonly available in those data sets, including age, sex, and study site. Similarly, we calculated DGE using predicted gene-expression profiles for frontal cortex obtained from *BrainGENIE* using the same mega-analysis approach. In addition, we inferred DGE signals for SCZ, BD, and ASD by applying the extension of *PrediXcan* compatible with GWAS summary statistics (*S-PrediXcan*) to published GWAS summary statistics for each disorder ^47–49^. Transcriptome-wide DGE effect sizes for each disorder obtained from peripheral blood mega-analyses *(t*-values), *BrainGENIE* mega-analyses (*t*-values), and *S-PrediXcan* (*z-*scores) were then compared with DGE effect sizes directly measured from *postmortem* brain published by the PsychENCODE Consortium (log_2_ fold-changes) using Pearson’s correlation test, which was chosen in order to assess the linear monotonic relationship between DGE signals derived from different methods.

### External Validation in the Religious Orders Study and Memory and Aging Project (ROSMAP)

To evaluate the generalizability of *BrainGENIE*’s imputation models, we leveraged a completely separate within-individual dataset (ROSMAP) that obtained paired measures of *ex vivo* peripheral blood (purified CD14+ monocytes) and *postmortem* brain transcriptome expression from bulk dorsolateral prefrontal cortex (DLPFC) and head of the caudate. The ROSMAP dataset included paired blood-DLPFC gene expression data for 109 older adults (age at first visit, range: 70.8 – 90+), including individuals diagnosed with Alzheimer’s disease (AD, *n*=54) or cognitively normal individuals (CN, *n*=55); in addition, we included paired blood-caudate gene expression data for 50 older adults diagnosed with AD (*n*=33) or CN (*n*=17) from a single sequencing batch (2×101bp on Illumina NovaSeq 6000). We restricted our analysis to individuals whose clinical diagnosis obtained at their last study visit was consistent with the final consensus diagnosis rendered at the time of death by expert review by one or more neurologists and a neuropsychologist. Standard preprocessing techniques were applied to the raw RNAseq data for each tissue, yielding normalized transcriptome expression levels for 14,639 genes in blood, 22,905 genes in DLPFC, and 16,380 genes in caudate. Detailed methods regarding our RNAseq pre-processing approaches are available in the Supplement. We evaluated two RNAseq normalization strategies on the RNAseq data from brain, which revealed that using residualized brain gene expression data (partialling out effects of *postmortem* interval, RIN, and batch) downwardly biased the accuracy and error of *BrainGENIE* (**Supplementary Figure 2**). The accuracy (Pearson’s *r*) of predicted and observed gene expression data was significantly higher for the normalized DLPFC RNAseq data compared with the residualized DLPFC RNAseq data (*t*=29.147, *df*=10400, *p*=1.3×10^−179^); this trend was also found for our analysis of the caudate (*t*=10.4, *df*=7163.5, *p*=4.2 × 10^−25^). Likewise, the mean square error of predicted and observed gene expression levels was significantly lower for the normalized DLPFC RNAseq data compared with the residualized DLPFC RNAseq data (*t*=-10.13, *df*=10932, *p*=5.2×10^−24^), which was consistent with the data from caudate (*t*=-3.27, *df*=7965.2, *p*=0.0011). Hence, we used the normalized brain RNAseq data instead of the residual brain RNAseq data (which were likely over-corrected) for our validation analyses. In addition, age and sex were not residualized from the RNAseq data from ROSMAP in order to evaluate the ability of *BrainGENIE* to recapitulate gene expression changes observed in *postmortem* brain related to those factors.

Imputation accuracy of *BrainGENIE* was measured using Pearson’s *r* comparing imputed brain gene expression to observed brain gene expression with both combined-sample and diagnostic group-specific evaluations. We performed linear regression analyses to measure changes in gene expression associated with diagnosis of AD (covarying for age and sex), then evaluated concordances for imputed brain *versus* observed brain transcriptome DGE signals, and between blood and brain DGE signals. Individuals in ROSMAP were much older than the donors in GTEx (v.8) whose data were used to train *BrainGENIE*. As such, a *post-hoc* exploratory analysis was performed to determine whether imputation performance was associated with aging among ROSMAP participants. Having identified an association of decreased imputation accuracy and age (**Supplementary Figure 3**), we performed a *post-hoc* analysis on individuals ≤ 85 years old among whom imputation accuracy was more reliable (DLPFC: AD=9, CN *n*=24; caudate: AD *n*=4, CN *n*=11).

### Concordance of Differential Gene Expression Profiles for Age and Sex

We evaluated how well *BrainGENIE* captures changes in gene expression associated with age and sex observed in the brain. Sex-specific and age-related effects have been linked with multiple brain disorders but their mechanisms are not well understood; thus, determining the effectiveness of *BrainGENIE* in modeling changes in gene expression related to these factors in the brain is warranted for future studies. We performed linear regression analyses in the CN group from ROSMAP to estimate the univariate effect size of age and sex on gene-expression levels observed in brain and blood, and *BrainGENIE*-imputed brain transcriptomes. We then used Pearson’s correlation coefficient to estimate the similarity of gene-expression changes related to sex and age between imputed brain *versus* observed brain transcriptomes, and between blood and brain transcriptomes.

## Results

### BrainGENIE Prediction Performance

**Table 1** summarizes the cross-validation prediction performance of *BrainGENIE* per brain region. **Table 2** depicts the number of genes that surpassed our criteria for being reliably predicted (*i*.*e*., average CV *R*^2^≥0.01, FDR*p*<0.05) per brain region, alongside the number of genes that *PrediXcan* was capable of imputing in the same regions, and the degree of overlap between *BrainGENIE* and *PrediXcan*. The statistics in **Table 1** and **Table 2** reflect the prediction performance for the top 20 PCs of blood-based transcriptome-wide gene expression, which yielded a higher average number of imputable genes per brain region relative to 5-, 10-, or 40-PC models. The prediction performance of *BrainGENIE*, measured by the average cross-validation *R*^2^, ranged from 0.08 – 0.16 for genes that met the criteria of reliable prediction by cross-validation (average CV *R*^2^≥0.01, FDR*p*<0.05). Brain tissues in GTEx showed overlap in the number of genes that were reliably predicted by *BrainGENIE* ranging from 303 (substantia nigra and anterior cingulate cortex) to 6,377 genes (putamen and frontal cortex [frozen]) (**Supplementary Figure 4A**). The proportion of overlapping genes reliably predicted by *BrainGENIE* in two brain tissues ranged from 17 – 69%. The pattern of overlap observed between brain tissues with respect to the number of genes reliably predicted by *BrainGENIE* mirrored the expected pattern of overlap between brain tissues with respect to the number of expressed genes among all GTEx donors (**Supplementary Figure 4B**), indicating that *BrainGENIE* preserves and recapitulates the spatial relationship between areas of the brain.

**Table 1.** Prediction accuracy of *BrainGENIE* (20-PC model) computed *via* 10-fold cross-validation of the GTEx Project (v.8) release across the 12 brain tissues.

| Brain Tissues | *N*<br>Samples | Mean Training $R^2$ | Mean CV $R^2$ | Max CV $R^2$ | *N* Reliably Predicted<br>Genes |
| --- | --- | --- | --- | --- | --- |
| Amygdala | 88 | 0.47 | 0.13 | 0.47 | 6,242 |
| Anterior cingulate cortex | 100 | 0.43 | 0.13 | 0.44 | 1,989 |
| Caudate | 138 | 0.35 | 0.09 | 0.39 | 6,421 |
| Cerebellum (Flash frozen) | 130 | 0.38 | 0.11 | 0.46 | 10,213 |
| Cerebellum (PAXgene fixed) | 153 | 0.37 | 0.10 | 0.47 | 7,437 |
| Frontal cortex (Flash frozen) | 125 | 0.34 | 0.10 | 0.57 | 11,569 |
| Frontal cortex (PAXgene fixed) | 142 | 0.34 | 0.10 | 0.41 | 3,603 |
| Hippocampus | 122 | 0.38 | 0.10 | 0.54 | 3,923 |
| Hypothalamus | 114 | 0.38 | 0.10 | 0.51 | 6,747 |
| Nucleus accumbens | 148 | 0.33 | 0.08 | 0.37 | 5,950 |
| Putamen | 125 | 0.40 | 0.11 | 0.42 | 9,797 |
| Substantia nigra | 86 | 0.51 | 0.16 | 0.67 | 1,733 |

Statistics are shown for genes that were considered reliably predicted based on the following criteria: cross-validation [CV] $R^2 \geq 0.01$ , CV FDRp $\leq 0.05$ .

**Table 2.**
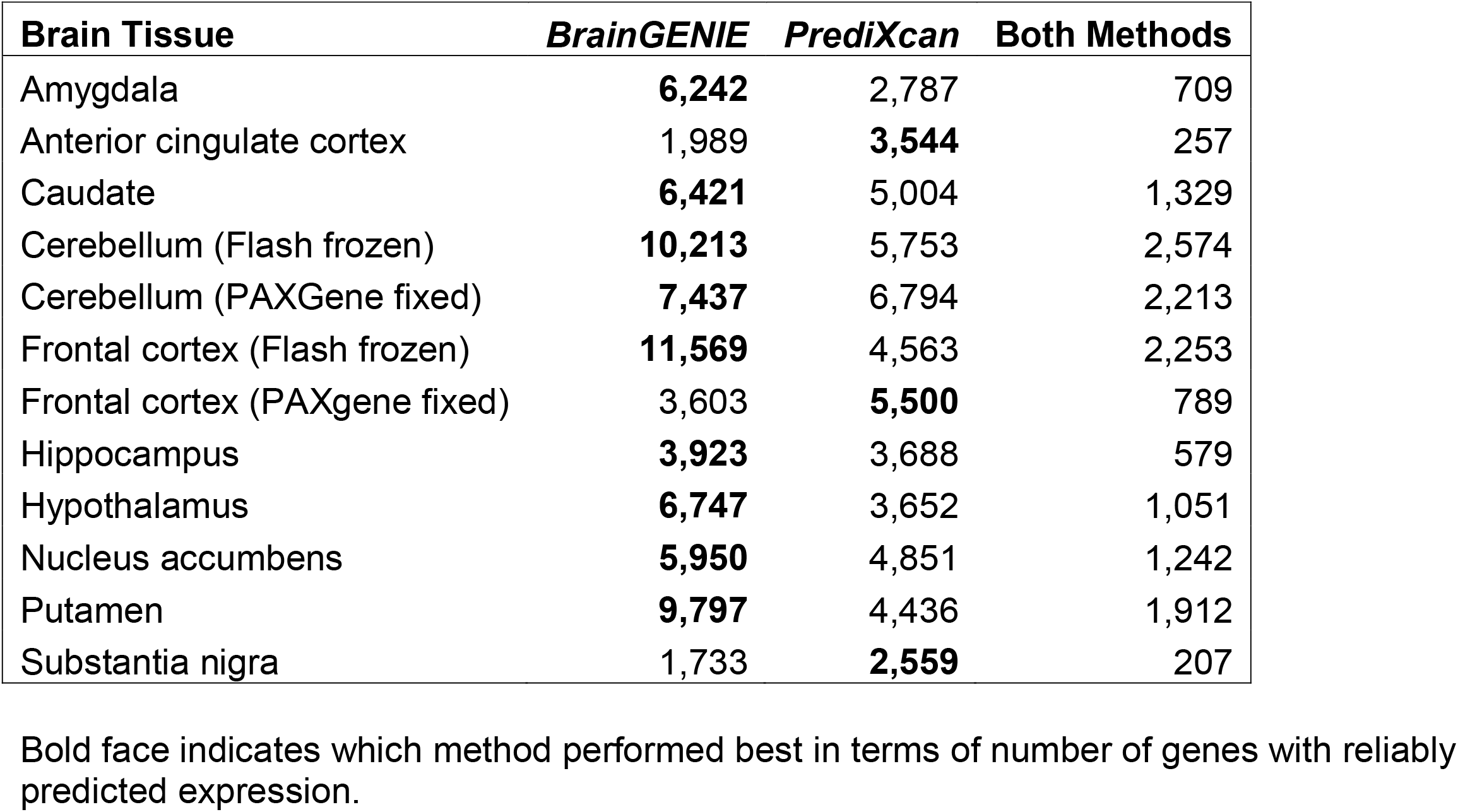
The number of genes for which brain-tissue-specific expression levels can be reliably predicted by *BrainGENIE* only (20-PC model), by *PrediXcan* only, or by both methods.

| Brain Tissue | *BrainGENIE* | *PrediXcan* | Both Methods |
| --- | --- | --- | --- |
| Amygdala | **6,242** | 2,787 | 709 |
| Anterior cingulate cortex | 1,989 | **3,544** | 257 |
| Caudate | **6,421** | 5,004 | 1,329 |
| Cerebellum (Flash frozen) | **10,213** | 5,753 | 2,574 |
| Cerebellum (PAXGene fixed) | **7,437** | 6,794 | 2,213 |
| Frontal cortex (Flash frozen) | **11,569** | 4,563 | 2,253 |
| Frontal cortex (PAXgene fixed) | 3,603 | **5,500** | 789 |
| Hippocampus | **3,923** | 3,688 | 579 |
| Hypothalamus | **6,747** | 3,652 | 1,051 |
| Nucleus accumbens | **5,950** | 4,851 | 1,242 |
| Putamen | **9,797** | 4,436 | 1,912 |
| Substantia nigra | 1,733 | **2,559** | 207 |

Bold face indicates which method performed best in terms of number of genes with reliably predicted expression.

The distributions of cross-validation *R*^2^ values produced by *BrainGENIE* and *PrediXcan* for all reliably predicted genes are shown in **Supplementary Figure 5**. The shapes of the distributions found using *BrainGENIE* were similar to those for *PrediXcan*; however, two distinctions were consistently noted across brain tissues. First, *PrediXcan* featured heavier right-tails compared to *BrainGENIE* (**Supplementary Figure 5**), indicative of *PrediXcan* having more genes with higher prediction accuracies. In contrast, *BrainGENIE* produced curves whose maxima were consistently shifted to the right relative to those of *PrediXcan*, indicative of greater average predictability with *BrainGENIE*. Disparities were seen in imputation performance for the pair of re-sampled brain tissues (frontal cortex and cerebellum), wherein the pair of brain tissues preserved in PAXgene tissue kits exhibited poorer imputation performance compared with the flash-frozen tissues. A likely explanation for this difference is that RNA qualities were better in the re-sampled flash-frozen tissues (**Supplementary Figure 6A**). The number of genes whose expression levels could be reliably predicted was strongly correlated with the average RNA quality of the brain tissues (Pearson’s *r*=0.71, *p*=0.0092, **Supplementary Figure 6B**).

The proportion of genes reliably predicted in the brain by *BrainGENIE* ranged from 9 – 57% of measured genes in the transcriptome (mean number of genes=6,302; range: 1,733 – 11,569 genes). The maximum average cross-validation prediction accuracy of *BrainGENIE* across all brain tissues ranged from *R*^2^=0.37 – 0.67. An average of 88% (range: 87.4 – 90%) of those genes whose expression levels were reliably predicted by *BrainGENIE* were not reliably predicted by *PrediXcan*. In contrast, an average of 73.2% of genes reliably predicted by *PrediXcan* were not reliably predicted by *BrainGENIE* (range: 50.6 – 92.3%). On average, 1,259 genes were found to be reliably predicted by both *BrainGENIE* and *PrediXcan* across 12 brain tissues (range: 207 [substantia nigra] – 2,574 genes [cerebellar hemisphere]).

Prediction accuracies generated by *BrainGENIE* were significantly better (two-tailed *t*-test FDR*p*<0.05) than those produced by *PrediXcan* for genes that both methods could reliably predict in one brain region: the substantia nigra (*n* genes=207, **Supplementary Table 2**). Among those genes that are reliably predicted by both methods, *PrediXcan* showed significantly better overall prediction accuracy for gene expression levels in nine brain tissues (**Supplementary Table 2**). Prediction accuracies were statistically indistinguishable between *BrainGENIE* and *PrediXcan* for the remaining two brain tissues: amygdala and anterior cingulate cortex (FDR*p*>0.05, **Supplementary Table 2**). When evaluating the similarity of prediction accuracy among genes that are reliably predicted by both methods, *BrainGENIE* showed a small negative correlation with *PrediXcan* for genes in frontal cortex (*r*=-0.08, *p*=1.6×10^−4^) and putamen (*r*=-0.05, *p*=0.046) (**Supplementary Table 3**), indicative of the methodological designs not converging to achieve consistent predictions.

### SynGO Gene-Set Enrichment

As shown in **Supplementary Table 4**, *BrainGENIE* reliably predicted expression levels of a significantly larger fraction of presynaptic and postsynaptic genes compared with *PrediXcan* for eight out of 12 brain tissues. *PrediXcan* predicted a larger fraction of presynaptic and postsynaptic genes than *BrainGENIE* in just three brain tissues: the anterior cingulate cortex, frontal cortex (PAXgene fixed), and substantia nigra. In the hippocampus, *BrainGENIE* predicted a larger proportion of postsynaptic (but not presynaptic) genes compared with *PrediXcan*. The largest difference between the two methods in terms of reliable prediction of pre- and post-synaptic genes was in frontal cortex, wherein the two methods showed an absolute difference of 57% (68% coverage by *BrainGENIE* versus 11% coverage by *PrediXcan*).

### Brain Cell-Type Marker-Gene Enrichment

*BrainGENIE* reliably predicted more human brain cell-type marker genes than *PrediXcan* for at least one brain cell type in 10 brain tissues (**Supplementary Table 5**). Markers for astrocytes, neurons, oligodendrocytes, and oligodendrocyte precursor cells (OPCs) were better predicted by *BrainGENIE* than by *PrediXcan* in six brain tissues; endothelial marker genes were better predicted by *BrainGENIE* in nine brain tissues; and microglia marker genes were better predicted by *BrainGENIE* in 10 brain tissues. The most significant difference that emerged in favor of *BrainGENIE* was the proportion of neuronal marker genes that were reliably predicted within the frontal cortex by *BrainGENIE* (65.2%) compared with *PrediXcan* (11.8%) (*χ*^2^=599.9, *p*=1.74×10^−132^, FDR*p*=1.25×10^−130^). *PrediXcan* better predicted neuronal markers in three brain tissues (anterior cingulate cortex, frontal cortex [PAXgene fixed], and substantia nigra), oligodendrocyte markers in two regions (anterior cingulate and substantia nigra), and OPC markers in cortex.

### Cross-Disorder Gene Sets Predicted by BrainGENIE versus PrediXcan

**Supplementary Figure 7** shows that, compared with *PrediXcan, BrainGENIE* reliably predicted more genes in 14 of the 45 Gene Ontology (GO) gene-sets found to have the strongest relationship with cross-disorder genetic risk for eight neuropsychiatric disorders.^23^ On average, *BrainGENIE* reliably predicted 1.73-fold (95% confidence interval [**CI**]: 1.62 – 1.84) more genes than *PrediXcan* across the 45 neuropsychiatric cross-disorder gene sets and 1.84-fold more genes than *PrediXcan* among the 14 significant gene sets. No cross-disorder gene sets were better predicted by *PrediXcan* than by *BrainGENIE*.

### Concordance of Differential Gene Expression Effect Sizes

Transcriptome-wide differential gene expression effect sizes for SCZ measured in peripheral blood were significantly correlated with differential gene expression effect sizes directly measured in *postmortem* brain from the PsychENCODE Consortium’s microarray meta-analysis (Pearson’s *r*=0.053, 95% CI: 0.035–0.072, *n* genes=10,832, *p*=2.6×10^−08^), and from the CommonMind Consortium (Pearson’s *r*=0.11, 95% CI: 0.093–0.133, *n* genes=9,434, *p*=2.1×10^−28^), but not the PsychENCODE Consortium’s RNAseq analysis (**Figure 1**). DGE effects for ASD from peripheral blood were inversely correlated with those from *postmortem* brain (**Figure 1**), which may reflect age differences between samples considering that individuals in the peripheral blood datasets were predominantly children whereas those in the *postmortem* brain studies were predominantly adults. DGE signals found using *S-PrediXcan* were not significantly correlated with *postmortem* brain DGE signals for SCZ, BD, or ASD (**Figure 1**). Conversely, DGE effects estimated from predicted genes’ expression profiles in brain using *BrainGENIE* were significantly correlated with results directly measured in *postmortem* brain (**Figure 1**). The strongest correlation that emerged was between DGE signals obtained using *BrainGENIE* and DGE signals directly measured in *postmortem* brain for SCZ found by the PsychENCODE Consortium’s microarray meta-analysis (Pearson’s *r*=0.52, 95% CI: 0.495–0.548, *n* genes=2,913, *p*=1.12×10^−203^). The DGE correlations for ASD, BD, and SCZ found between *BrainGENIE* and *postmortem* brain showed significant replication in an independent PsychENCODE Consortium cohort profiled *via* RNAseq (**Figure 1**). Furthermore, the SCZ DGE correlation between *BrainGENIE* and *postmortem* brain showed significant replication in a second independent cohort from the CommonMind Consortium (Pearson’s *r*=0.34, 95% CI: 0.33–0.37, *n* genes=8,077, *p* = 4.9×10^−230^, **Figure 1**). The DGE agreement between *BrainGENIE*-imputed brain gene expression and *postmortem* brain measured gene expression was significantly stronger than between peripheral blood and *postmortem* brain for ASD, BD, and SCZ for at least one of the four *BrainGENIE* models (*z*-test *p*-values<0.05, **Supplementary Table 6**). Similarly, measured DGE signals from *postmortem* brain were significantly more concordant with those predicted from *BrainGENIE* than from *S-PrediXcan* for ASD, BD, and SCZ (**Supplementary Table 6**).

**Figure 1.**
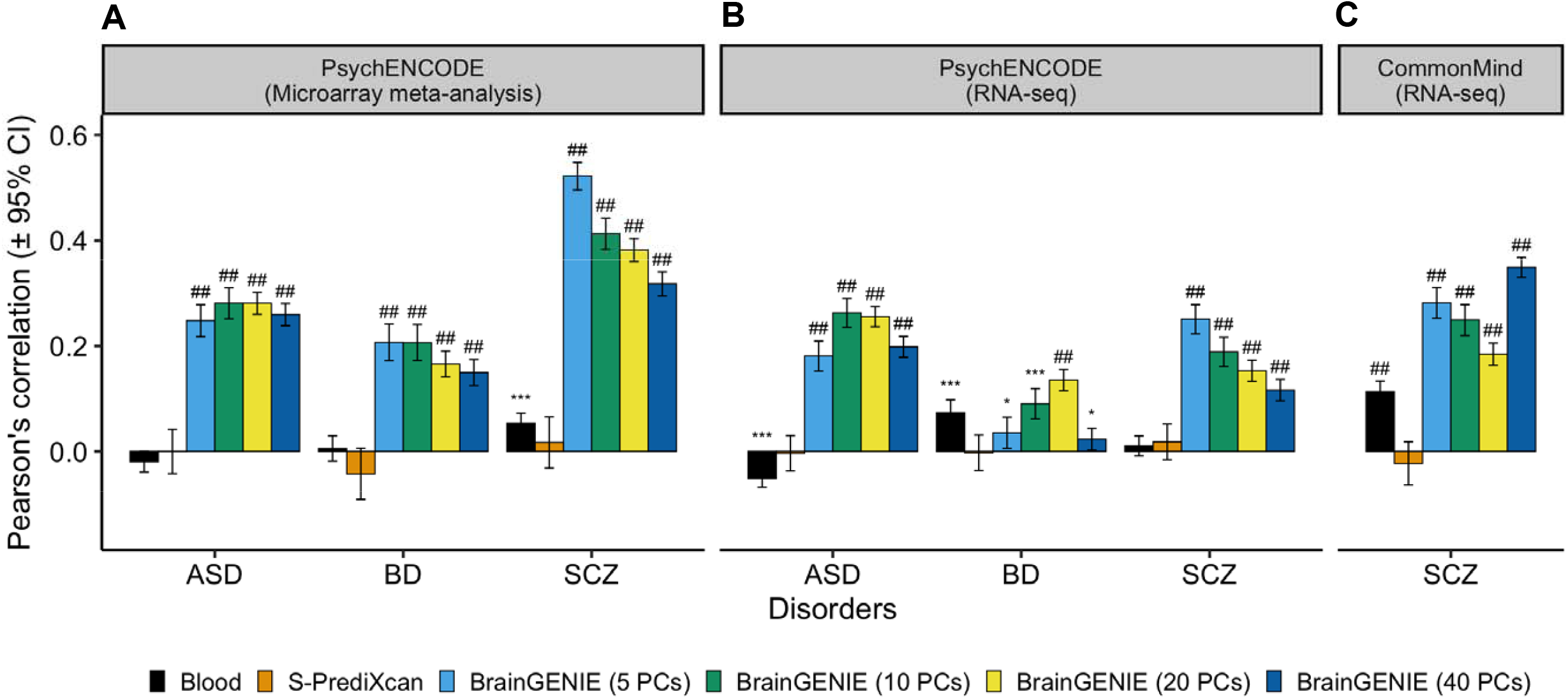
Concordance of case-control differential gene expression (DGE) signals obtained by *BrainGENIE* and *S-PrediXcan* compared to (**A**) DGE signals derived from *postmortem* cortical microarray meta-analyses for ASD, BD, and SCZ, (**B**) DGE signals derived from RNA-sequencing analysis for ASD, BD, and SCZ by the PsychENCODE Consortium, and (**C**) DGE signals obtained from *postmortem* prefrontal cortex RNA-sequencing analysis for SCZ by the CommonMind Consortium. Abbreviations: autism spectrum disorder (ASD), bipolar disorder (BD), and schizophrenia (SCZ). Symbols for significance thresholds: *p*<0.05 (*), FDR*p*≤1×10^−5^ (***), FDR*p*≤1×10^−10^ (#), FDR*p*≤1×10^−20^ (##).

### Validation of BrainGENIE’s DLPFC Models in ROSMAP

Among all genes that were predicted by *BrainGENIE* in ROSMAP for the DLPFC (*n*=5,955), 2,581 genes (43%) were validated in terms of predicted gene-expression levels being concordant with observed gene-expression levels (*R*^2^≥0.01). When we stratified by diagnosis, a larger number of genes showed concordance between predicted and observed DLPFC expression levels in the CN group (3,343, 56%) than in AD cases (1,924, 32%) (**Supplementary Figure 8**); we also found that DLPFC prediction accuracies were significantly higher in the CN group relative to AD cases (*t*=25.49, *df*=11853, *p*=1.5×10^−139^). We found a positive correlation between imputation accuracy of *BrainGENIE* and average DLPFC expression levels of genes in GTEx (v.8) in each diagnostic group (**Supplementary Figure 9**), indicating that genes expressed at higher levels in the DLPFC were more reliably predicted.

We correlated gene expression levels between blood and brain to evaluate the magnitude of cross-tissue concordance, and whether measured brain transcriptomes were better predicted by measured blood gene expression or by *BrainGENIE*-imputed brain gene expression. Among 3,078 genes that were found to be expressed in both blood and DLPFC samples from ROSMAP, Pearson’s *r-*values were significantly lower when correlating brain and blood gene expression levels (mean *r*=0.016) than when correlating brain and *BrainGENIE*-imputed gene expression (mean *r*=0.069; two-tailed *t*=-18.7, *df*=6755.2, *p*=3.8×10^−76^). In our analysis of all CN individuals from ROSMAP, *BrainGENIE* predictions and *postmortem* brain measurements showed similar effect sizes for genes that differed by sex (*r*=0.22, *p*<2.2×10^−16^) and age (*r*=0.15, *p*<2.2×10^−16^) (**Supplementary Figure 10A**). Unexpectedly, diagnostic-group DGE effect-sizes for AD diagnosis estimated in brain-imputed transcriptomes were significantly negatively correlated with effect-sizes directly measured in *postmortem* brain transcriptomes (*r*=-0.074, *p*=2.2×10^−9^) (**Supplementary Figure 10A**). Our *post-hoc* inspection revealed that imputation performance was diminished among ROSMAP donors at extremely old age (>85 years old, **Supplementary Figure 3**), which potentially explained the negative correlations observed for DGE effect-sizes for AD diagnosis in DLPFC. The concordance of DGE effect-sizes in the DLPFC for sex (*r*=0.43, *p*<2.2×10^−16^), age (*r*=0.34, *p*<2.2×10^−16^), and AD diagnosis (*r*=0.50, *p*<2.2×10^−16^) were strongly and significantly positive when we restricted correlation analysis of *BrainGENIE* and *postmortem* DLPFC results to donors in ROSMAP aged ≤ 85 years old (**Figure 2A**). Hence, individuals at extreme old age may introduce error in the predicted gene-expression levels in the full ROSMAP sample, leading to downwardly biased results from our correlation analyses. DGE effect-sizes estimated in peripheral blood transcriptomes showed a relatively weak concordance with effect-sizes directly measured in *postmortem* brain for age, sex, and AD in the full ROSMAP sample *and* in individuals ≤ 85 years old for DLPFC (**Supplementary Figure 10B** and **Figure 10B**). This finding was consistent with the patterns from our DGE-concordance analyses of ASD, BD, and SCZ (**Figure 1**), suggesting that *BrainGENIE* yields more reliable estimates of brain-specific transcriptomic changes than direct blood-to-brain correlations.

**Figure 2.**
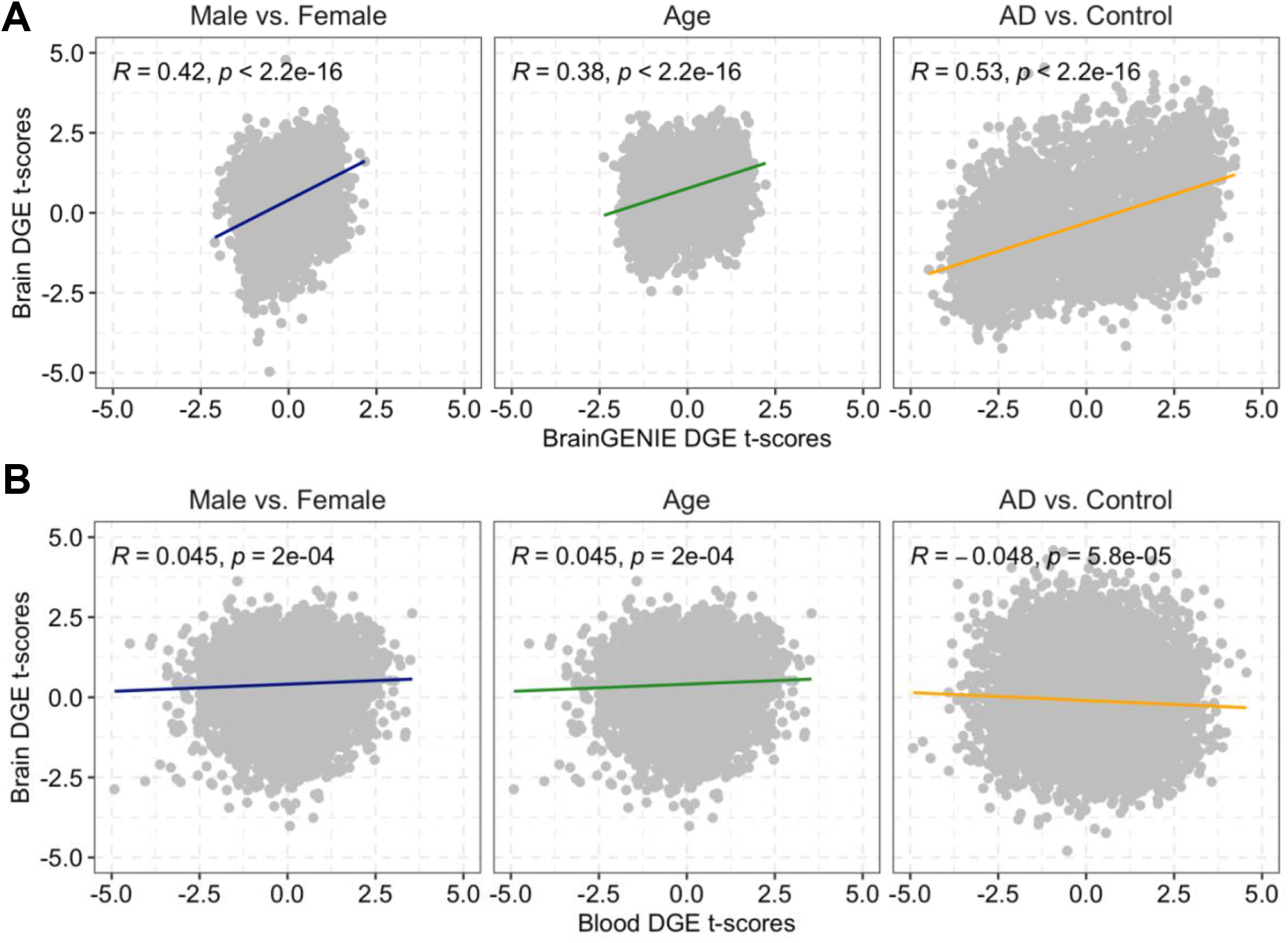
Correlation analysis of differential gene expression (DGE) effect-sizes for age, sex, and Alzheimer’s disease (AD) obtained from ROSMAP subjects aged ≤ 85 years old with paired blood and DLPFC transcriptomes (*n*=33, AD *n*=9, Control *n*=24). Scatterplots show the paired samples correlation of differential gene expression effect-sizes to evaluate concordance of (**A**) frontal cortex-imputed and observed DLPFC transcriptomes, and (**B**) CD14+ purified monocyte and DLPFC transcriptomes.

### Validation of BrainGENIE’s Caudate Models in ROSMAP

Among the 3,265 genes that were imputed in the caudate, 747 genes (23%) were validated in terms of predicted gene-expression levels being concordant with observed gene-expression levels. Prediction accuracies of gene-expression levels in the caudate were higher in AD cases compared with CN controls (*t*=8.67, *df*=6235.7, *p*=5.3×10^−18^). When stratified by diagnosis, predicted expression levels of 910 genes (28%) in AD cases and 859 genes (26%) in CN controls were concordant with observed gene-expression levels in caudate. In contrast to DLPFC, a negative correlation between imputation accuracy and average caudate expression levels of genes in CN controls and AD cases emerged, indicating that genes expressed at higher levels in the caudate were less reliably predicted (**Supplementary Figure 9**). DGE effect-sizes were significantly positively correlated across *BrainGENIE*-imputed results compared with *postmortem* caudate results for sex (*r*=0.42, *p*<2.2×10^−16^) and AD diagnosis (*r*=0.15, p<2.2×10^−16^), but not age (*r*=-0.015, *p*=0.4) (**Supplementary Figure 11A**). Restricting our analysis to individuals ≤ 85 years old resulted in a stronger correlation of DGE effect-sizes for sex (*r*=0.57, *p*<2.2×10^−16^), age (*r*=0.093, *p*<4.8×10^−9^), and AD diagnosis (r=0.48, *p*<2.2×10^−16^) across *BrainGENIE*-imputed results compared with *postmortem* caudate (**Figure 3A**). DGE results derived from peripheral blood for age, sex, and AD diagnosis showed less similarity with results derived from *postmortem* caudate in the full ROSMAP sample *and* individuals ≤ 85 years old (**Supplementary 11B** and **Figure 3B**).

**Figure 3.**
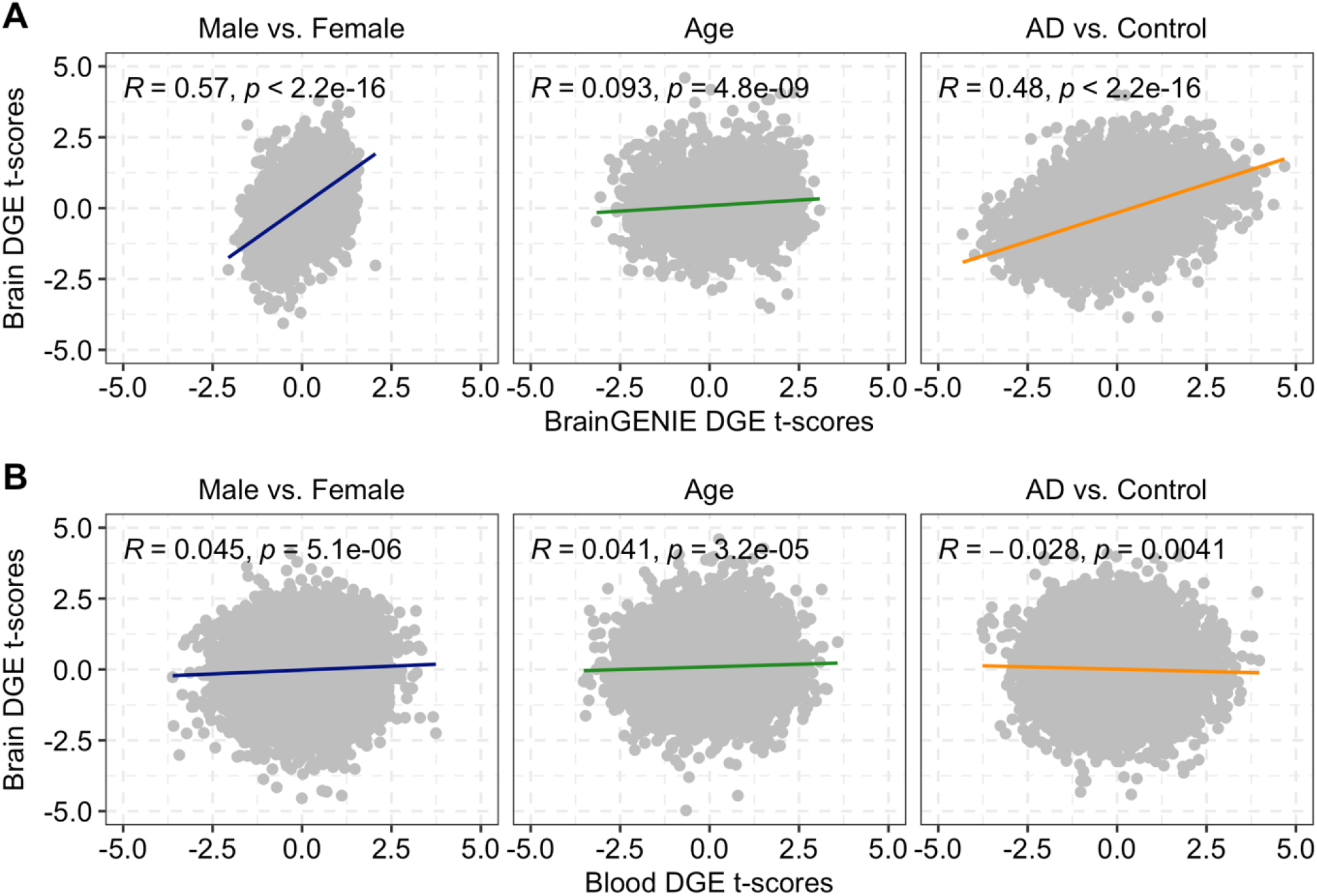
Correlation analysis of differential gene expression (DGE) effect-sizes for age, sex, and Alzheimer’s disease (AD) obtained from ROSMAP subjects aged ≤ 85 years old with paired blood and caudate transcriptomes (*n*=15, AD *n*=4, Control *n*=11). Scatterplots show the paired samples correlation of differential gene expression effect-sizes to evaluate concordance of (**A**) caudate-imputed and observed brain transcriptomes, and (**B**) CD14+ purified monocyte and caudate transcriptomes.

## Discussion

This study introduced and benchmarked a novel computational method called *BrainGENIE*, which predicts brain-tissue-specific gene expression levels based on peripheral blood transcriptomes. Over the past decade, there has been rapid growth in the number of blood-based transcriptome studies aimed at identifying biomarkers for neuropsychiatric disorders. This has led to a vast amount of useful data that may hold untapped information about the brain. Much of the raw data from published blood-based transcriptome studies of neuropsychiatric disorders can be readily downloaded from public repositories (*i*.*e*., Gene Expression Omnibus [GEO], ArrayExpress) or made available to investigators with controlled access (*i*.*e*., dbGaP, NIMHGR, Synapse). It is therefore possible to further mine those stores of transcriptome data with *BrainGENIE*, thus generating novel mechanistic hypotheses about disease and advancing our understanding of brain disorders in a way that is clearly superior to direct correlation of blood and brain measures.

*BrainGENIE* exploits PCA as an efficient method for dimensionality-reduction while capturing more variability in the blood transcriptome for prediction compared to individual gene transcripts. PCA also helps to reduce the potential of overfitting; *i*.*e*., imposing a limit on the number of input features steers a model away from “learning” random noise and failing to generalize to external data. However, the risk for over-fitting is not eliminated by PCA alone. We applied a standard machine learning approach of *k*-fold cross-validation to estimate the ability of *BrainGENIE* to generalize its predictions on unseen data, and then performed the vitally important external validation of our models (for frontal cortex and caudate) in a fully independent ROSMAP cohort. Further validation is warranted to determine the generalizability of our prediction models for other brain tissues and with an external dataset that is more closely matched to the demographic and technical parameters observed in our training set. In the future, modifications to *BrainGENIE* may allow for subsets of genes to be best predicted by PCs and others by individual gene transcripts, or collections of a few (or more) closely correlated transcripts.

The predictive performance of *BrainGENIE* was affected by a number of factors, including the number of samples available for training and quality of the extracted RNAs. Brain tissues that had large sample sizes (*i*.*e*., frontal cortex) showed better prediction performance relative to brain regions with fewer samples (*i*.*e*., amygdala, substantia nigra). In addition, imputation performance improved with RNA quality of the brain tissues. Differences in imputation performance were seen between pairs of re-sampled tissue collected from frontal cortex and cerebellum. The pair of tissues collected and preserved in PAXgene fixative exhibited lower RIN values (likely due to degraded RNAs) and lower imputation performance than the pair of brain tissues that were shipped to University of Miami Endowment Brain Bank for collection and preservation by flash freezing. Given the differences observed between flash frozen and fixed brain tissues, we recommend that investigators prioritize *BrainGENIE* imputation results derived from flash frozen brain tissues. Furthermore, there were on average 48 fewer donors (range: 28 – 63) with paired blood-brain transcriptome data in GTEx (the required input for *BrainGENIE*) than donors with paired genetic and transcriptome data (the required input for *PrediXcan*). Given this disparity in sample size, it is challenging to draw strong conclusions about differences in model performance for individual genes between *BrainGENIE* and *PrediXcan*, but generally *PrediXcan* would be in a more powerful position to outperform *BrainGENIE*. This was not generally observed however, and in fact *BrainGENIE* outperformed *PrediXcan* on a number of benchmarks. Thus, global differences between the methods could not be explained by variation in sample size alone, and even as *BrainGENIE* was limited by a smaller sample size for model-training, we found that *BrainGENIE* could impute a substantial fraction of genes that were not imputable using *PrediXcan*. This suggests that non-genetic components of gene expression ignored by *PrediXcan* models hold reliably useful information for transcriptome imputation.

GTEx’s transcriptomic data were derived from bulk *postmortem* brain tissue; thus, we did not model gene expression for any specific brain-cell type. We instead modeled the cross-tissue overlap at the level of cell mixtures in the brain and blood for *BrainGENIE*. It is possible that commonalities seen between brain and blood gene expression could be driven by a possible shared lineage between macrophages and microglia ^52,53^. Nevertheless, studies have implicated glial cell dysfunction in neuropsychiatric disorders ^54^, which makes them an interesting target for gene-expression modeling. The specific neuronal component of brain gene-expression levels, either directly measured in *postmortem* brain or as imputed by *BrainGENIE*, is unknown; however, we did show that brain-cell-specific markers were capable of being reliably predicted by *BrainGENIE*, indicating that cell-type-specific modeling may be feasible. Furthermore, inferences may be made about how different brain-cell types explain components of gene expression through cell deconvolution analysis, which can be accomplished by several algorithms that make use of gene-expression data from single cells.

The current version of *BrainGENIE* can predict expression levels for 1,733 to 11,569 genes in the human brain (depending on the brain region), which accounts for about 9 to 57% of the brain transcriptome. Prior iterations of *BrainGENIE* have made continual improvements in the number of reliably predicted genes and the variance accounted for in brain-tissue-specific gene-expression levels by moving from an approach that used individual gene transcripts in blood to predict brain gene-expression levels to the current approach of using principal components of blood gene expression. This suggests that further refinement of our models will continue to improve predictions until they reach their (unknown) maximum per gene and per brain region. As one would expect, not all or even most genes are imputable with *BrainGENIE*, but the number of *new* genes that can be imputed with *BrainGENIE* and not by *PrediXcan* is considerable. The amount of overlap between *BrainGENIE* and *PrediXcan* in terms of genes whose expression can be reliably predicted was relatively small. In addition, prediction accuracies were not well correlated between *BrainGENIE* and *PrediXcan*, indicating that the different modeling approaches achieve partially orthogonal outcomes when predicting brain transcriptomes. This suggests that there is value to integrating *BrainGENIE* and *PrediXcan* for a combined and complementary approach to transcriptome imputation wherever genotypes and blood gene-expression data are available from the same individuals. Ideally, the strengths of multiple modeling approaches like those in *BrainGENIE, PrediXcan*, and others, would be combined into a unified framework (or through the integration of outputs from multiple disparate models) to deliver a holistic portrait of the landscape of the human brain transcriptome.

We suggest the following tool selection depending on the type of data available for transcriptome imputation: 1) if only transcriptome data are available from blood, use *BrainGENIE*, 2) if only GWAS data are available, use *PrediXcan* (or a derivative), 3) if blood transcriptome *and* GWAS data are available, use *BrainGENIE* and *PrediXcan* (or a derivative) to achieve best predicted expression levels on a *per-*gene basis depending on the target brain region. For genes that are predictable by both methods, use the method that achieved better accuracy for the specific gene being imputed in the target brain region.

Our results showed that transcriptome-wide DGE effect sizes observed directly in *postmortem* brain were in better agreement with DGE effect sizes predicted using *BrainGENIE* than with DGE effect sizes found in analyses of peripheral blood and those imputed by *PrediXcan*. This advantage of *BrainGENIE* over peripheral blood and *PrediXcan* was most striking for SCZ but was still evident for BD and ASD. Likewise, in our within-individual validation analysis of the ROSMAP dataset, DGE effect sizes for AD exhibited stronger agreement between *BrainGENIE* and brain than between peripheral blood and brain. Concordance of DGE effect sizes between *BrainGENIE* and *postmortem* brain varied based on the number of PCs included in the imputation models. This finding may encourage investigators to parameterize the number of PCs for *BrainGENIE* based on the model that yields best overall prediction accuracy. However, it is important to consider which genes are included (or lost) or more reliably predicted when adjusting the number of PCs used by *BrainGENIE*, as this can be relevant for downstream analyses. For example, a study focused on frontal-cortical expression of the SCZ risk-gene complement component 4 (*C4A*) would favor the 20-PC model (average CV *R*^2^=0.20, *p*=3.8×10^−7^) as it yielded higher accuracy than the *BrainGENIE* models with 5-, 10-, and 40-PCs. Alternatively, an example whereby the 40-PC model yields better imputation is for hippocampal expression of presenilin-1 (*PSEN1*), a leading risk factor for the early-onset form of Alzheimer’s disease (CV *R*^2^=0.07, *p*=0.0039).

An interesting finding that emerged for ASD from our DGE concordance analysis was the inverse correlation between DGE measured in peripheral blood and DGE directly measured in *postmortem* brain. There are a variety of possible explanations for this finding, but no definitive conclusion could be drawn here. One consideration is that the participants in the ASD dataset are mostly young children and toddlers (mean age=5.8 years, 95% CI: 4.8 – 5.3), whereas the GTEx dataset used to train *BrainGENIE* contained only adults (aged 20 – 70). Additionally, the PsychENCODE dataset for ASD contained mostly adults (mean age=29.7 years, 95% CI: 26.3 – 33.1). Nevertheless, these results suggest that *BrainGENIE* needs to be further optimized on individuals out-of-range relative to the age of the GTEx (or other emergent) training samples. Extreme old age, in particular, was found to affect *BrainGENIE* performance in ROSMAP. The ages of some ROSMAP samples were out-of-range compared to our GTEx training data, though we were able to mitigate the detrimental effect of age on imputation performance by excluding individuals over 85 years old.

Among the retained ROSMAP samples, we demonstrated that *BrainGENIE* generalized well to an external sample, wherein performance was best for the unaffected CN group. A larger number of reliably predicted genes, and higher imputation accuracies, were obtained in the CN group relative to AD cases; hence, it is possible that brain pathology partly disrupts the blood-to-brain mappings expected by the imputation models. Disruption of brain-imputed gene expression accuracy may itself become a biomarker for disease, a prospect to be evaluated more fully later. Despite such factors that affect imputation performance, the brain transcriptomes imputed by *BrainGENIE* outperformed peripheral blood gene expression as a proxy for brain-transcriptome profiles. Specifically, we found that transcriptomic signatures of sex, age, and AD measured in brain samples from ROSMAP were more consistent with those imputed by *BrainGENIE* than those measured in peripheral blood. Taken together, *BrainGENIE* provides a more accurate modeling of the brain compared with peripheral blood. While it is evident that subsets of peripheral blood -omic features show moderate-to-strong overlap with - omic profiles in the brain ^9^, we showed that measured blood gene expression is not as strong a surrogate as *BrainGENIE*-imputed gene expression for transcriptomic measures of the *postmortem* brain.

We applied statistical corrections to remove effects of age, sex, and genetic ancestry from the gene-expression data so that those factors would not systematically bias our models. Still, it is possible that characteristics of the GTEx sample are not fully representative of the entire population. For example, donors in the GTEx Project were predominantly of European ancestry, hence limiting applicability of transcriptome-imputation across diverse ancestral groups. Amassing large sample sizes that encompass a broader range of characteristics (*e*.*g*., environmental exposures, genetic background, and demographics, to name a few) would allow *BrainGENIE* to make use of more biological (useful) variability that may help increase the number of reliably predicted genes and improve variance accounted for. Increasing sample ascertainment from diverse human populations, coupled with deeper phenotyping, are strategic ways to enable more effective transcriptome-imputation modelling.

In sum, *BrainGENIE* is a novel and validated approach to investigating brain-tissue-specific gene-expression profiles. We demonstrated that gene-expression changes associated with disease and imputed in the brain by *BrainGENIE* were in better agreement (relative to *cis-* eQTL-based predictions of gene expression by *PrediXcan* and to gene-expression changes detected in peripheral blood) with corresponding gene-expression changes detected in studies of *postmortem* brain. The main challenge of transcriptome-imputation is identifying a model and set of predictor variables that can efficiently and reliably predict gene-expression levels while ensuring that downstream analyses of predicted expression levels can yield biologically meaningful results. *PrediXcan* and *FUSION*, respectively, can reliably predict an average of 18% and 16% of the brain transcriptome (compared with an average of 30% by *BrainGENIE*). Those methods have been successful in identifying novel tissue-specific dysregulation of gene expression in complex disorders. A strength of *BrainGENIE* is that it can capture reliable impacts of non-genetic factors of gene expression regulation that are not yet modeled by *cis*-eQTL-based methods. *BrainGENIE* fills a void in the study of the brain transcriptome by both allowing analyses of genes that were not previously imputable and improving the predictability of disease-relevant gene sets that *PrediXcan* can only partially impute.

Though we showed that *BrainGENIE* has advantages over conceptually similar methods, our intention is for it to serve as a complement to genetic-based transcriptome-imputations methods. In practice, our recommendation would be to integrate *BrainGENIE* with other methods whenever possible, to boost confidence in gene-disease associations, hence permitting a deeper understanding of complex phenotypes. As such, *BrainGENIE* offers an important function in systems-level research into the brain and serves as a valuable hypothesis-generating tool for mechanistic studies. Potential applications of *BrainGENIE* are far-reaching and would be best suited (relative to *PrediXcan* and *FUSION*) to study gene expression longitudinally, including: across developmental timepoints of the brain, pre- and post-exposure (*e*.*g*., environmental risks, traumatic life experiences), and modeling the effects of medication or other clinical interventions. *BrainGENIE* also could be used to impute brain-tissue-specific transcriptomes at any point in a person’s lifetime, opening the possibility that we could find causal and longitudinal mechanisms underlying neuropsychiatric disease. The reach of our toolset can be extended with additional developments to achieve reliable imputations of cell-type specific transcriptomes, and transcriptomes of other inaccessible tissues, as well as models of alternatively spliced mRNAs and short and long noncoding RNAs, all of which are feasible objectives.

## Data and code availability

Data and source code can be accessed from the following GitHub repository: https://github.com/hessJ/BrainGENIE. This software was developed and tested on a laptop running MacOS Big Sur (2.6GHz 6-Core Intel i7 processor, 16 GB 2400 MHz DDR4). *BrainGENIE* has linear running time based on the input sample size (runtime for one brain region: 4.5 seconds for 1,000 samples, 30.7 seconds for 10,000 samples).

## Appendix

^†^ Alphabetical list of authors in the Neuropsychiatric Consortium for the Analysis and Sharing of Transcriptomes (NCAST)

**Natalie Jane Beveridge**; Schizophrenia Research Institute, Sydney, New South Wales, Australia; School of Biomedical Sciences & Pharmacy, College of Health, Medicine and Wellbeing, The University of Newcastle, New South Wales, Australia; Centre for Brain & Mental Health, University of Newcastle, Callaghan, Newcastle, Australia

**Vaughan Carr**; Schizophrenia Research Institute, Sydney, New South Wales, Australia; School of Psychiatry, University of New South Wales, Kensington, New South Wales, Australia

**Simone de Jong**; MRC Social, Genetic and Developmental Psychiatry Centre, King’s College London, London, UK.

**Erin Gardiner**; Schizophrenia Research Institute, Sydney, New South Wales, Australia; School of Biomedical Sciences & Pharmacy, College of Health, Medicine and Wellbeing, The University of Newcastle, New South Wales, Australia; Centre for Brain & Mental Health, University of Newcastle, Callaghan, Newcastle, Australia

**Brian Kelly**; School of Medicine & Public Health, The University of Newcastle, Callaghan, Newcastle, Australia; Hunter Medical Research Institute, Newcastle, Australia; Centre for Brain & Mental Health, University of Newcastle, Callaghan, Newcastle, Australia

**Nishantha Kumarasinghe**; School of Medicine & Public Health, The University of Newcastle, Callaghan, Newcastle, Australia; Department of Anatomy, Faculty of Medical Sciences, University of Sri Jayawardenepura, Nugegoda, Sri Lanka; Schizophrenia Research Institute, Sydney, New South Wales, Australia; Faculty of Medicine, Sir John Kotelawala Defence University, Ratmalana, Sri Lanka

**Roel A. Ophoff**; Center for Neurobehavioral Genetics, Semel Institute for Neuroscience and Human Behavior, University of California Los Angeles, Los Angeles, CA, USA; Department of Human Genetics, University of California, Los Angeles, CA, USA.

**Ulrich Schall**; School of Medicine & Public Health, The University of Newcastle, Callaghan, Newcastle, Australia; Schizophrenia Research Institute, Sydney, New South Wales, Australia; Hunter Medical Research Institute, Newcastle, Australia; Centre for Brain & Mental Health, University of Newcastle, Callaghan, Newcastle, Australia

**Rodney J. Scott**; School of Biomedical Sciences & Pharmacy, College of Health, Medicine and Wellbeing, The University of Newcastle, New South Wales, Australia; Hunter Medical Research Institute, Newcastle, Australia

**Boryana Stamova**; Department of Neurology, UC Davis School of Medicine, Sacramento, California

**Paul Tooney**; Schizophrenia Research Institute, Sydney, New South Wales, Australia; School of Biomedical Sciences & Pharmacy, College of Health, Medicine and Wellbeing, The University of Newcastle, New South Wales, Australia; Hunter Medical Research Institute, Newcastle, Australia; Centre for Brain & Mental Health, University of Newcastle, Callaghan, Newcastle, Australia

## Acknowledgements

The Genotype-Tissue Expression (GTEx) Project was supported by the Common Fund of the Office of the Director of the National Institutes of Health (commonfund.nih.gov/GTEx). Additional funds were provided by the NCI, NHGRI, NHLBI, NIDA, NIMH, and NINDS. Donors were enrolled at Biospecimen Source Sites funded by NCI\Leidos Biomedical Research, Inc. subcontracts to the National Disease Research Interchange (10XS170), Roswell Park Cancer Institute (10XS171), and Science Care, Inc. (X10S172). The Laboratory, Data Analysis, and Coordinating Center (LDACC) was funded through a contract (HHSN268201000029C) to the The Broad Institute, Inc. Biorepository operations were funded through a Leidos Biomedical Research, Inc. subcontract to Van Andel Research Institute (10ST1035). Additional data repository and project management were provided by Leidos Biomedical Research, Inc.(HHSN261200800001E). The Brain Bank was supported supplements to University of Miami grant DA006227. Statistical Methods development grants were made to the University of Geneva (MH090941 & MH101814), the University of Chicago (MH090951,MH090937, MH101825, & MH101820), the University of North Carolina - Chapel Hill (MH090936), North Carolina State University (MH101819),Harvard University (MH090948), Stanford University (MH101782), Washington University (MH101810), and to the University of Pennsylvania (MH101822). The results published here are in whole or in part based on data obtained from the AD Knowledge Portal (https://adknowledgeportal.org). Study data were provided by the Rush Alzheimer’s Disease Center, Rush University Medical Center, Chicago. Data collection was supported through funding by NIA grants P30AG10161 (ROS), R01AG15819 (ROSMAP; genomics and RNAseq), R01AG17917 (MAP), R01AG30146, R01AG36042 (5hC methylation, ATACseq), RC2AG036547 (H3K9Ac), R01AG36836 (RNAseq), R01AG48015 (monocyte RNAseq) RF1AG57473 (single nucleus RNAseq), U01AG32984 (genomic and whole exome sequencing), U01AG46152 (ROSMAP AMP-AD, targeted proteomics), U01AG46161(TMT proteomics), U01AG61356 (whole genome sequencing, targeted proteomics, ROSMAP AMP-AD), the Illinois Department of Public Health (ROSMAP), and the Translational Genomics Research Institute (genomic). Additional phenotypic data can be requested at www.radc.rush.edu. The datasets used for the analyses described in this manuscript were obtained from dbGaP at http://www.ncbi.nlm.nih.gov/gap through dbGaP accession number phs000424.v8.p2. Dr. Hess is supported by grants from the U.S. National Institute of Mental Health (R21MH126494-01) and NARSAD: The Brain & Behavior Research Foundation. Dr. Kong is supported by grants R01MH107205 and R24OD024622. Dr. Faraone is supported by the European Union’s Seventh Framework Programme for research, technological development and demonstration under grant agreement no 602805, the European Union’s Horizon 2020 research and innovation programme under grant agreements No 667302 & 728018 and NIMH grants 5R01MH101519, U01 MH109536-01, and R21MH126494-01. Dr. Glatt is supported by grants from the U.S. National Institutes of Health (R01MH101519, R01AG054002, R01AG064955, and R21MH126494-01), the Sidney R. Baer, Jr. Foundation, and NARSAD: The Brain & Behavior Research Foundation.

