## Supplemental Information for "*BrainGENIE*: The Brain Gene Expression and Network Imputation Engine"

### Abbreviations used

| **Term** | **Abbreviated** |
|---|---|
| Oligodendrocyte precursor cells | OPCs |
| False discovery rate adjusted *p*-value | FDRp |
| Standard deviation | s.d. |
| Dorsolateral prefrontal cortex | DLPFC |

### Supplementary Methods

#### *Data Import and Quality Control of Array Data*

We applied a standardized pre-processing pipeline for microarray data that we previously developed to convert raw probe intensities into a suitable format for meta- and mega-analysis of multiple studies [1]. The following steps were applied to unprocessed microarray data sets acquired from our literature search: (1) conventional or GC-corrected robust multi-array averaging (RMA) performed on a study-wise basis to exon array and gene chips from Affymetrix, respectively (using the *R* package *affy* [2]), (2) adjusting probe-level data for background intensities based on negative control probes available on Illumina microarrays (using the *R* package *limma* [3]), (3) $\log_2$ transformation to stabilize variance, and (4) quantile normalization to minimize between-sample variation. Microarray probes were mapped to HGNC gene symbols. Probes that did not map to a known a HGNC gene symbol were discarded as these have unknown biological relevance. Sometimes a gene was matched to multiple probes, in which case we computed the median normalized expression, thus converting probe-level microarray data to gene-level expression values. Gene expression values were z-transformed (mean=0, s.d.=1) *per* study to minimize unwanted variation due to platform differences. Standardized gene expression data were merged across studies based on common gene symbols.

### Monocyte RNA-sequencing Data from ROSMAP

All data from monocyte RNA-sequences were analyzed by the uniform procedures described below. FASTQ files were downloaded from Synapse (https://www.synapse.org/#!Synapse:syn3388564) for two batches of sequenced reads generated by a Illumina HiSeq2500 sequencer. The cDNA libraries for these samples were prepared from 20-2000 FACS-sorted monocytes using a SMART-seq2 protocol. Sequencing quality was accessed by fastQC v0.11.8 program, and multiple quality statistics were collected and evaluated for each sample, including per base/tile sequence quality, adapter content, per sequence GC content, duplication level report and etc. Then low-quality bases/reads and adapters were trimmed by trimmomatic v0.39 with a quality score cutoff of 30. Next, the trimmed reads were mapped to Gencode GRCh38.p13 release 35 Human reference genome using STAR aligner v2.7.3a. Mapping statistics were collected from alignment results and percentages of mapping to genome regions were summarized using PicardTools v2.21.6. Then reads mapped to genes were summarized by featureCounts program in subread v1.6.4. Raw count data for each sample are combined and annotated to reference genome using customized R scripts.

After carefully examining multiple quality matrix and visualizing data by MDS plots, Low quality samples and outliers were removed from further analyses. Genes including protein-coding genes, miRNAs, lncRNAs were kept with CPM ≥ 1 in at least 50% samples. Counts with 0s were replaced with 0.1, and data were normalized to effective library size by TMM method in edgeR v3.28.1 program.

### *Postmortem* Brain RNA-sequencing Data from ROSMAP

A matrix of preprocessed gene transcript quantifications derived from bulk *postmortem* dorsolateral prefrontal cortex (DLPFC) tissue samples collected by ROSMAP and sequenced on a Illumina HiSeq 2000 (101x2 bp) were downloaded from the Synapse database (https://www.synapse.org/#!Synapse:syn3388564). Gene transcript quantifications were expressed as fragments per kilobase per million mapped reads (FPKM). We discarded gene transcripts with no or exceptionally low expression levels (FPKM < 0.1 in > 10 samples), then transformed FPKM values to $\log_2$ scale. A many-to-one relationship of multiple Ensembl transcript IDs mapping to the same HGNC gene ID was resolved by collapsing transcript cluster expression levels by computing the median. We downloaded BAM files that stored aligned RNA-sequencing data from the *postmortem* head of caudate sequenced by Illumina NovaSeq 6000. BAM files were converted to FASTQ files using the software *Picard* (v.2.21.6). Sequencing quality was assessed by *fastQC* (v.0.11.8) and *multiQC* (v1.8). Low quality bases were removed from raw reads using *Trimmomatic* (v.0.39). Trimmed reads were aligned to the human genome (GRCh38.p13.r35.pri) using *STAR* (v.2.73a). Reads mapped to genes (protein-coding genes, miRNAs, and lncRNAs) were quantified by *featureCounts* package in *subread* (v.1.6.4). Genes with low expression (< 1 counts per million [CPM] for 50% of samples) were discarded. Gene counts were scaled to $\log_2$CPM then were normalized by the TMM method from the *R* package *edgeR*.

### Supplementary Tables

#### Supplementary Table 1. Description of peripheral blood case-control datasets included in our differential gene expression mega-analyses.

| Study ID | Platform | # of cases | # of controls | % male | Mean age in years (s.d.) |
|---|---|---|---|---|---|
| **Schizophrenia datasets** | | | | | |
| Gardiner et al. (2013) [6] | Illumina Human HT-12 v3 BeadChip | 78 | 78 | 51.3 | 39.5 (13.0) |
| Glatt et al. (2009) [7] | Affymetrix Human Exon 1.0 ST | 13 | 8 | 66.7 | 44.1 (7.8) |
| Glatt et al. (2011) [8] | Affymetrix HG U133 Plus 2.0 | 8 | 12 | 50 | 40.8 (11.6) |
| Kumarasinghe et al. (2013) [9] | Illumina Human HT-12 v3 BeadChip | 9 | 11 | 60 | 34.5 (13.0) |
| de Jong et al. (2012) [10] | Illumina Human Ref-8 v3.0 | 15 | 21 | 72.2 | 30.1 (11.3) |
| de Jong et al. (2012) [10] | Illumina Human HT-12 v3 BeadChip | 106 | 95 | 58.7 | 39.5 (12.5) |
| Tsuang et al. (2005) [11] | Affymetric HG U133A/Affymetrix HG U133 Plus 2.0 | 29 | 16 | 46.7 | 35.7 (9.8) |
| ***Total*** | | **258** | **241** | | |
| **Bipolar Disorder datasets** | | | | | |
| **Study ID** | **Platform** | **# of cases** | **# of controls** | **% male** | **Mean age in years (s.d.)** |
| Beech et al. (2010) [12] | Illumina Human-6 v2 BeadChip | 20 | 15 | 31.4 | 34.1 (10.5) |
| Bousman et al. (2010) [13] | Affymetrix Human Exon 1.0 ST | 9 | 8 | 70.6 | 43.6 (7.2) |
| Clelland et al. (2013) [14] | Affymetrix HG U133 Plus 2.0 | 26 | 25 | 100 | 36.6 (13.0) |
| Padmos et al. (2008) [15] | Affymetrix U95v2 | 5 | 6 | 45.5 | 23.6 (9.6) |
| Savitz et al. (2013) [16] | Illumina HT-12 v4 expression BeadChip | 8 | 24 | 46.9 | 35.0 (11.1) |
| Tsuang et al. (2005)[11] | Affymetric HG U133A/Affymetrix HG U133 Plus 2.0 | 16 | 23 | 41 | 42.4 (13.5) |
| ***Total*** | | **84** | **101** | | |
| **Autism datasets** | | | | | |
| **Study ID** | **Platform** | **# of cases** | **# of controls** | **% male** | **Mean age in years (s.d.)** |
| CHARGE[17–20] | Affymetrix HG U133 Plus 2.0 | 118 | 90 | 85.6 | 3.7 (0.76) |
| Glatt et al. (2012)[21] | Illumina HumanWG-6 v3.0 | 173 | 159 | 69.2 | 2.0 (0.78) |
| Kong et al. (2012)[22] | Affymetrix HG U133 Plus 2.0/Affymetrix Human Exon 1.0 ST | 165 | 106 | 80.4 | 8.1 (4.2) |
| Alter et al. (2011) [23] | Affymetrix HG U133 Plus 2.0 | 75 | 59 | 100 | 6.6 (2.4) |
| Kong et al. (2013)[24] | Affymetrix Human Exon 1.0 ST | 53 | 17 | 80 | 9.6 (3.8) |
| ***Total*** | | **584** | **431** | | |

Supplementary Table 2. Summary of two-tailed *t*-tests comparing the prediction accuracy for genes between BrainGENIE and PrediXcan. The analysis was restricted to genes that both methods can reliably predict.

| Brain Tissue | # of genes | BrainGENIE (avg. Pearson's *r*) | PrediXcan (avg. Pearson's *r*) | *t*-statistic | *p*-value | 95% CI Lower | 95% CI Upper | FDRp |
|---|---|---|---|---|---|---|---|---|
| Amygdala | 709 | 0.36 | 0.35 | 1.86 | 0.06 | -0.001 | 0.021 | 0.07 |
| Anterior cingulate cortex BA24 | 257 | 0.36 | 0.35 | 0.56 | 0.58 | -0.013 | 0.024 | 0.58 |
| Caudate basal ganglia | 1329 | 0.29 | 0.34 | -10.00 | 5.49E-23 | -0.052 | -0.035 | 1.32E-22 |
| Cerebellum (Flash frozen) | 2574 | 0.32 | 0.38 | -16.90 | 6.40E-62 | -0.063 | -0.050 | 7.68E-61 |
| Cerebellum (PAXgene fixed) | 2213 | 0.31 | 0.37 | -16.07 | 5.73E-56 | -0.067 | -0.053 | 3.44E-55 |
| Frontal cortex (Flash frozen) | 2253 | 0.31 | 0.35 | -11.74 | 2.70E-31 | -0.046 | -0.033 | 1.08E-30 |
| Frontal cortex (PAXgene fixed) | 789 | 0.31 | 0.34 | -5.40 | 8.42E-08 | -0.041 | -0.019 | 1.39E-07 |
| Hippocampus | 579 | 0.31 | 0.34 | -5.39 | 9.27E-08 | -0.046 | -0.021 | 1.39E-07 |
| Hypothalamus | 1051 | 0.30 | 0.33 | -6.39 | 2.24E-10 | -0.037 | -0.020 | 4.47E-10 |
| Nucleus accumbens basal ganglia | 1242 | 0.28 | 0.33 | -11.01 | 3.02E-27 | -0.057 | -0.040 | 9.06E-27 |
| Putamen basal ganglia | 1912 | 0.32 | 0.34 | -4.81 | 1.62E-06 | -0.024 | -0.010 | 2.16E-06 |
| Substantia nigra | 207 | 0.40 | 0.37 | 2.96 | 3.32E-03 | 0.010 | 0.050 | 3.98E-03 |

Supplementary Table 3. Summary statistics from Pearson's correlation tests evaluating the similarity of prediction accuracies between *BrainGENIE* and *PrediXcan* for genes that both methods could reliably predict.

| **Brain tissue** | **Pearson's *r*** | **95% CI Lower** | **95% CI Upper** | **z-statistic** | **p-value** |
|---|---|---|---|---|---|
| Amygdala | -0.07 | -0.140 | 0.007 | -1.78 | 0.08 |
| Anterior cingulate cortex BA24 | -0.08 | -0.198 | 0.045 | -1.25 | 0.21 |
| Caudate basal ganglia | 0.00 | -0.053 | 0.055 | 0.04 | 0.97 |
| Cerebellum (Flash frozen) | -0.04 | -0.075 | 0.002 | -1.87 | 0.06 |
| Cerebellum (PAXgene fixed) | 0.00 | -0.047 | 0.037 | -0.23 | 0.82 |
| Frontal cortex (Flash frozen) | -0.08 | -0.120 | -0.038 | -3.77 | 1.6e-4 |
| Frontal cortex (PAXgene fixed) | 0.01 | -0.064 | 0.075 | 0.16 | 0.88 |
| Hippocampus | -0.04 | -0.124 | 0.038 | -1.04 | 0.30 |
| Hypothalamus | -0.02 | -0.084 | 0.037 | -0.77 | 0.44 |
| Nucleus accumbens basal ganglia | -0.01 | -0.063 | 0.048 | -0.26 | 0.79 |
| Putamen basal ganglia | -0.05 | -0.090 | -0.001 | -2.00 | 0.046 |
| Substantia nigra | -0.05 | -0.183 | 0.089 | -0.69 | 0.49 |

Supplementary Table 4. Two-tailed Chi-square tests of equality comparing the proportion of presynaptic and postsynaptic gene sets reliably predicted by *BrainGENIE* relative to *PrediXcan*. Results highlighted in red denote that *BrainGENIE* predicted a significantly larger fraction of synaptic gene sets compared to *PrediXcan*.

| SynGO: Pre-synaptic genes (k=747) | | | | | | 95% Confidence Interval | |
|---|---|---|---|---|---|---|---|
| **Brain tissue** | **% reliably predicted by BrainGENIE** | **% reliably predicted by PrediXcan** | $\chi^2$ | ***p*-value** | **FDRp** | **Lower** | **Upper** |
| Amygdala | 32 | 7.6 | 138 | 7.19E-32 | 2.16E-31 | 0.2 | 0.28 |
| Anterior cingulate cortex BA24 | 8.7 | 9.5 | 0.2 | 6.53E-01 | 6.53E-01 | -0.04 | 0.02 |
| Caudate basal ganglia | 20.5 | 15.4 | 6.2 | 1.26E-02 | 1.89E-02 | 0.01 | 0.09 |
| Cerebellum (Flash frozen) | 55 | 22 | 171.1 | 4.27E-39 | 1.71E-38 | 0.28 | 0.38 |
| Cerebellum (PAXgene fixed) | 39.8 | 27 | 26.6 | 2.52E-07 | 4.32E-07 | 0.08 | 0.18 |
| Frontal cortex (Flash frozen) | 68.5 | 12.2 | 490.5 | 1.10E-108 | 1.32E-107 | 0.52 | 0.61 |
| Frontal cortex (PAXgene fixed) | 12 | 13.9 | 1 | 3.17E-01 | 3.46E-01 | -0.05 | 0.02 |
| Hippocampus | 13.5 | 11.2 | 1.6 | 2.09E-01 | 2.51E-01 | -0.01 | 0.06 |
| Hypothalamus | 28 | 8.2 | 97.7 | 4.90E-23 | 9.80E-23 | 0.16 | 0.24 |
| Nucleus accumbens basal ganglia | 35.1 | 11.4 | 116.3 | 4.14E-27 | 9.93E-27 | 0.19 | 0.28 |
| Putamen basal ganglia | 50.5 | 13.4 | 234.6 | 5.92E-53 | 3.55E-52 | 0.33 | 0.42 |
| Substantia nigra | 4.1 | 6.4 | 3.4 | 6.44E-02 | 8.58E-02 | -0.05 | 0 |
| SynGO: Post-synaptic genes (k=1,351) | | | | | | 95% Confidence Interval | |
| **Brain region** | **% reliably predicted by BrainGENIE** | **% reliably predicted by PrediXcan** | $\chi^2$ | ***p*-value** | **FDRp** | **Lower** | **Upper** |
| Amygdala | 33.2 | 5.3 | 337.2 | 2.65E-75 | 1.06E-74 | 0.25 | 0.31 |
| Anterior cingulate cortex BA24 | 8.2 | 9.9 | 2.2 | 1.40E-01 | 1.53E-01 | -0.04 | 0.01 |
| Caudate basal ganglia | 22.1 | 14.4 | 26.8 | 2.22E-07 | 3.80E-07 | 0.05 | 0.11 |
| Cerebellum (Flash frozen) | 53.8 | 21.7 | 295.3 | 3.52E-66 | 1.05E-65 | 0.29 | 0.36 |
| Cerebellum (PAXgene fixed) | 36.3 | 28 | 20.9 | 4.81E-06 | 7.21E-06 | 0.05 | 0.12 |
| Frontal cortex (Flash frozen) | 67.3 | 10.8 | 902.9 | 2.28E-198 | 2.73E-197 | 0.53 | 0.6 |
| Frontal cortex (PAXgene fixed) | 15.5 | 15 | 0.1 | 7.48E-01 | 7.48E-01 | -0.02 | 0.03 |
| Hippocampus | 14.6 | 9.8 | 13.7 | 2.14E-04 | 2.86E-04 | 0.02 | 0.07 |
| Hypothalamus | 28.3 | 7.5 | 198.7 | 3.93E-45 | 9.43E-45 | 0.18 | 0.24 |
| Nucleus accumbens basal ganglia | 34.8 | 12.2 | 190.2 | 2.81E-43 | 5.62E-43 | 0.19 | 0.26 |
| Putamen basal ganglia | 58.8 | 13.8 | 589.9 | 2.59E-130 | 1.55E-129 | 0.42 | 0.48 |
| Substantia nigra | 4.6 | 6.3 | 3.5 | 6.20E-02 | 7.44E-02 | -0.03 | 0 |

Supplementary Table 5. Two-proportions Chi-square test of equality reveals that *BrainGENIE* predicted a larger proportion of human brain cell-type marker gene sets compared to *PrediXcan*. Results highlighted in red denote that *BrainGENIE* predicted a significantly larger fraction of cell-type marker genes compared to *PrediXcan*.

| Brain tissue | Cell type | % reliably predicted by BrainGENIE | % reliably predicted by PrediXcan | $\chi^2$ | *p*-value | FDRp | 95% Confidence interval Lower | 95% Confidence interval Upper |
|---|---|---|---|---|---|---|---|---|
| Amygdala | Astrocytes | 21.9 | 9 | 62.7 | 2.39E-15 | 7.49E-15 | 0.10 | 0.16 |
| Amygdala | Endothelial | 24.4 | 7.4 | 106.8 | 4.94E-25 | 2.73E-24 | 0.14 | 0.20 |
| Amygdala | Microglia | 22.4 | 4.6 | 134.1 | 5.08E-31 | 3.33E-30 | 0.15 | 0.21 |
| Amygdala | Neuron | 41.6 | 6.3 | 340.1 | 5.96E-76 | 1.07E-74 | 0.32 | 0.39 |
| Amygdala | Oligodendrocyte | 12.7 | 9.4 | 5.2 | 2.25E-02 | 3.11E-02 | 0.00 | 0.06 |
| Amygdala | OPCs | 16.6 | 7.6 | 18.2 | 1.99E-05 | 3.11E-05 | 0.05 | 0.13 |
| Anterior cingulate cortex BA24 | Astrocytes | 6.2 | 12.3 | 21.4 | 3.65E-06 | 6.25E-06 | -0.09 | -0.03 |
| Anterior cingulate cortex BA24 | Endothelial | 9.4 | 9.4 | 0 | 1.00E+00 | 1.00E+00 | -0.03 | 0.03 |
| Anterior cingulate cortex BA24 | Microglia | 4.9 | 5.8 | 0.6 | 4.27E-01 | 4.92E-01 | -0.03 | 0.01 |
| Anterior cingulate cortex BA24 | Neuron | 2.8 | 9 | 33.5 | 7.09E-09 | 1.59E-08 | -0.08 | -0.04 |
| Anterior cingulate cortex BA24 | Oligodendrocyte | 4 | 10.3 | 29 | 7.42E-08 | 1.48E-07 | -0.09 | -0.04 |
| Anterior cingulate cortex BA24 | OPCs | 8 | 8.8 | 0.1 | 7.32E-01 | 7.99E-01 | -0.04 | 0.03 |
| Caudate basal ganglia | Astrocytes | 20 | 19.5 | 0.1 | 8.22E-01 | 8.84E-01 | -0.03 | 0.04 |
| Caudate basal ganglia | Endothelial | 26.7 | 16 | 33.5 | 7.29E-09 | 1.59E-08 | 0.07 | 0.14 |
| Caudate basal ganglia | Microglia | 23.8 | 11.3 | 53.1 | 3.12E-13 | 8.64E-13 | 0.09 | 0.16 |
| Caudate basal ganglia | Neuron | 13.7 | 14 | 0 | 8.97E-01 | 9.36E-01 | -0.03 | 0.03 |
| Caudate basal ganglia | Oligodendrocyte | 29.1 | 18.1 | 32.9 | 9.47E-09 | 2.01E-08 | 0.07 | 0.15 |
| Caudate basal ganglia | OPCs | 22.2 | 13.8 | 11.4 | 7.39E-04 | 1.11E-03 | 0.03 | 0.13 |
| Cerebellum (Flash frozen) | Astrocytes | 29.9 | 20 | 25.6 | 4.10E-07 | 7.58E-07 | 0.06 | 0.14 |
| Cerebellum (Flash frozen) | Endothelial | 36 | 17.9 | 82.3 | 1.18E-19 | 4.71E-19 | 0.14 | 0.22 |
| Cerebellum (Flash frozen) | Microglia | 25.4 | 12.5 | 53.3 | 2.81E-13 | 8.09E-13 | 0.09 | 0.16 |
| Cerebellum (Flash frozen) | Neuron | 41 | 21 | 92.6 | 6.50E-22 | 2.93E-21 | 0.16 | 0.24 |
| Cerebellum (Flash frozen) | Oligodendrocyte | 37.3 | 19.7 | 75.1 | 4.38E-18 | 1.66E-17 | 0.14 | 0.22 |
| Cerebellum (Flash frozen) | OPCs | 30.8 | 15.2 | 33.5 | 7.21E-09 | 1.59E-08 | 0.10 | 0.21 |
| Cerebellum (PAXgene fixed) | Astrocytes | 27 | 24.9 | 1 | 3.08E-01 | 3.69E-01 | -0.02 | 0.06 |
| Cerebellum (PAXgene fixed) | Endothelial | 28.2 | 19.6 | 19.9 | 8.32E-06 | 1.36E-05 | 0.05 | 0.12 |
| Cerebellum (PAXgene fixed) | Microglia | 23.2 | 15.4 | 19 | 1.28E-05 | 2.06E-05 | 0.04 | 0.11 |
| Cerebellum (PAXgene fixed) | Neuron | 28.2 | 25.4 | 1.9 | 1.73E-01 | 2.22E-01 | -0.01 | 0.07 |
| Cerebellum (PAXgene fixed) | Oligodendrocyte | 26.5 | 22.7 | 3.7 | 5.47E-02 | 7.43E-02 | 0.00 | 0.08 |
| Cerebellum (PAXgene fixed) | OPCs | 23 | 19.4 | 1.7 | 1.88E-01 | 2.38E-01 | -0.02 | 0.09 |
| Frontal cortex (Flash frozen) | Astrocytes | 24.3 | 15.7 | 22.6 | 2.02E-06 | 3.54E-06 | 0.05 | 0.12 |
| Frontal cortex (Flash frozen) | Endothelial | 57.6 | 12.4 | 447 | 3.19E-99 | 1.15E-97 | 0.41 | 0.49 |
| Frontal cortex (Flash frozen) | Microglia | 44.9 | 8.9 | 327.7 | 3.03E-73 | 4.37E-72 | 0.32 | 0.40 |
| Frontal cortex (Flash frozen) | Neuron | 65.2 | 11.8 | 599.9 | 1.75E-132 | 1.26E-130 | 0.50 | 0.57 |
| Frontal cortex (Flash frozen) | Oligodendrocyte | 32.1 | 14 | 91.3 | 1.21E-21 | 5.14E-21 | 0.14 | 0.22 |
| Frontal cortex (Flash frozen) | OPCs | 35.4 | 12 | 74.4 | 6.34E-18 | 2.28E-17 | 0.18 | 0.29 |
| Frontal cortex (PAXgene fixed) | Astrocytes | 5.6 | 21.5 | 106.6 | 5.57E-25 | 2.86E-24 | -0.19 | -0.13 |
| Frontal cortex (PAXgene fixed) | Endothelial | 17.4 | 17.8 | 0 | 8.60E-01 | 9.11E-01 | -0.04 | 0.03 |
| Frontal cortex (PAXgene fixed) | Microglia | 10.5 | 12.2 | 1.3 | 2.59E-01 | 3.22E-01 | -0.05 | 0.01 |
| Frontal cortex (PAXgene fixed) | Neuron | 4.9 | 15.8 | 62.9 | 2.23E-15 | 7.29E-15 | -0.14 | -0.08 |
| Frontal cortex (PAXgene fixed) | Oligodendrocyte | 13.8 | 16.6 | 2.8 | 9.26E-02 | 1.24E-01 | -0.06 | 0.00 |
| Frontal cortex (PAXgene fixed) | OPCs | 8.6 | 14.4 | 7.7 | 5.51E-03 | 7.94E-03 | -0.10 | -0.02 |
| Hippocampus | Astrocytes | 14 | 12.7 | 0.6 | 4.30E-01 | 4.92E-01 | -0.02 | 0.04 |
| Hippocampus | Endothelial | 20.5 | 10.2 | 40 | 2.50E-10 | 6.42E-10 | 0.07 | 0.14 |
| Hippocampus | Microglia | 14.2 | 6.8 | 28.4 | 1.01E-07 | 1.97E-07 | 0.05 | 0.10 |
| Hippocampus | Neuron | 4.6 | 7.3 | 6 | 1.40E-02 | 1.97E-02 | -0.05 | -0.01 |
| Hippocampus | Oligodendrocyte | 15.4 | 15.4 | 0 | 1.00E+00 | 1.00E+00 | -0.03 | 0.03 |
| Hippocampus | OPCs | 12.4 | 10.8 | 0.5 | 4.89E-01 | 5.51E-01 | -0.03 | 0.06 |
| Hypothalamus | Astrocytes | 24.3 | 13.3 | 38.9 | 4.43E-10 | 1.10E-09 | 0.08 | 0.14 |
| Hypothalamus | Endothelial | 29.9 | 8.8 | 141.3 | 1.39E-32 | 9.99E-32 | 0.18 | 0.25 |
| Hypothalamus | Microglia | 22.9 | 6.5 | 105.9 | 7.58E-25 | 3.64E-24 | 0.13 | 0.20 |
| Hypothalamus | Neuron | 25.6 | 7.6 | 115.7 | 5.48E-27 | 3.29E-26 | 0.15 | 0.21 |

| | | | | | | | | |
|---|---|---|---|---|---|---|---|---|
| Hypothalamus | Oligodendrocyte | 11.6 | 11.8 | 0 | 9.45E-01 | 9.72E-01 | -0.03 | 0.03 |
| Hypothalamus | OPCs | 22.8 | 9.8 | 30 | 4.27E-08 | 8.79E-08 | 0.08 | 0.18 |
| Nucleus accumbens basal ganglia | Astrocytes | 30.8 | 18.9 | 37.3 | 1.02E-09 | 2.46E-09 | 0.08 | 0.16 |
| Nucleus accumbens basal ganglia | Endothelial | 31.1 | 15.1 | 71.2 | 3.30E-17 | 1.13E-16 | 0.12 | 0.20 |
| Nucleus accumbens basal ganglia | Microglia | 21.8 | 9.4 | 57.5 | 3.46E-14 | 1.04E-13 | 0.09 | 0.16 |
| Nucleus accumbens basal ganglia | Neuron | 19.1 | 11.6 | 21.1 | 4.42E-06 | 7.41E-06 | 0.04 | 0.11 |
| Nucleus accumbens basal ganglia | Oligodendrocyte | 21.7 | 16.3 | 9.1 | 2.52E-03 | 3.70E-03 | 0.02 | 0.09 |
| Nucleus accumbens basal ganglia | OPCs | 16.4 | 13.8 | 1.1 | 2.89E-01 | 3.53E-01 | -0.02 | 0.07 |
| Putamen basal ganglia | Astrocytes | 44.5 | 16.5 | 183.6 | 7.90E-42 | 7.11E-41 | 0.24 | 0.32 |
| Putamen basal ganglia | Endothelial | 53.6 | 12.9 | 371.3 | 9.53E-83 | 2.29E-81 | 0.37 | 0.45 |
| Putamen basal ganglia | Microglia | 29.7 | 8.1 | 150.8 | 1.17E-34 | 9.33E-34 | 0.18 | 0.25 |
| Putamen basal ganglia | Neuron | 48.1 | 11.7 | 314.3 | 2.48E-70 | 2.98E-69 | 0.33 | 0.40 |
| Putamen basal ganglia | Oligodendrocyte | 49.7 | 19.2 | 204.6 | 2.05E-46 | 2.10E-45 | 0.26 | 0.35 |
| Putamen basal ganglia | OPCs | 29.8 | 11.4 | 50.6 | 1.12E-12 | 2.98E-12 | 0.13 | 0.23 |
| Substantia nigra | Astrocytes | 9.2 | 7.4 | 1.9 | 1.68E-01 | 2.20E-01 | -0.01 | 0.04 |
| Substantia nigra | Endothelial | 5.9 | 7.1 | 1 | 3.18E-01 | 3.76E-01 | -0.03 | 0.01 |
| Substantia nigra | Microglia | 11.9 | 5.5 | 25 | 5.78E-07 | 1.04E-06 | 0.04 | 0.09 |
| Substantia nigra | Neuron | 1.8 | 4.8 | 13.2 | 2.83E-04 | 4.34E-04 | -0.05 | -0.01 |
| Substantia nigra | Oligodendrocyte | 3.6 | 9.5 | 27.5 | 1.59E-07 | 3.01E-07 | -0.08 | -0.04 |
| Substantia nigra | OPCs | 7 | 5.8 | 0.4 | 5.18E-01 | 5.74E-01 | -0.02 | 0.04 |

Supplementary Table 6. Statistical test for differences in concordance of differential gene expression signals obtained from *BrainGENIE*, peripheral blood, and *S-PrediXcan* relative to independent *postmortem* brain cohorts (PsychENCODE, CommonMind). Results that remained significant after multiple testing correction appear in bold.

| Concordance with differential gene expression signals from PsychENCODE (Microarray meta-analysis) | | | | | | |
|---|---|---|---|---|---|---|
| **Disorder** | **Method 1** | **Method 2** | **Δ Pearson's *r*** | ***z*-value** | ***p*-value** | **FDRp** |
| ASD | Blood | BrainGENIE (20 PCs) | -0.30 | -20.21 | 3.68E-91 | **9.67E-90** |
| ASD | Blood | BrainGENIE (40 PCs) | -0.28 | -18.73 | 1.49E-78 | **2.60E-77** |
| ASD | Blood | BrainGENIE (10 PCs) | -0.30 | -16.14 | 6.35E-59 | **5.13E-58** |
| ASD | Blood | BrainGENIE (5 PCs) | -0.27 | -14.14 | 1.04E-45 | **7.78E-45** |
| ASD | BrainGENIE (20 PCs) | S-PrediXcan | 0.28 | 11.95 | 3.12E-33 | **1.56E-32** |
| ASD | BrainGENIE (40 PCs) | S-PrediXcan | 0.26 | 11.01 | 1.73E-28 | **7.00E-28** |
| ASD | BrainGENIE (10 PCs) | S-PrediXcan | 0.28 | 10.80 | 1.75E-27 | **6.79E-27** |
| ASD | BrainGENIE (5 PCs) | S-PrediXcan | 0.25 | 9.43 | 2.11E-21 | **6.71E-21** |
| ASD | BrainGENIE (5 PCs) | BrainGENIE (20 PCs) | -0.03 | -1.75 | 3.97E-02 | **4.85E-02** |
| ASD | BrainGENIE (5 PCs) | BrainGENIE (10 PCs) | -0.03 | -1.53 | 6.33E-02 | 7.31E-02 |
| ASD | BrainGENIE (20 PCs) | BrainGENIE (40 PCs) | 0.02 | 1.41 | 7.89E-02 | 9.00E-02 |
| ASD | BrainGENIE (10 PCs) | BrainGENIE (40 PCs) | 0.02 | 1.16 | 1.23E-01 | 1.36E-01 |
| ASD | Blood | S-PrediXcan | -0.02 | -0.81 | 2.08E-01 | 2.20E-01 |
| ASD | BrainGENIE (5 PCs) | BrainGENIE (40 PCs) | -0.01 | -0.61 | 2.70E-01 | 2.81E-01 |
| ASD | BrainGENIE (10 PCs) | BrainGENIE (20 PCs) | 0.00 | 0.01 | 4.98E-01 | 4.98E-01 |
| BD | Blood | BrainGENIE (10 PCs) | -0.20 | -9.29 | 7.60E-21 | **2.35E-20** |
| BD | Blood | BrainGENIE (5 PCs) | -0.20 | -9.20 | 1.83E-20 | **5.34E-20** |
| BD | Blood | BrainGENIE (20 PCs) | -0.16 | -9.08 | 5.20E-20 | **1.47E-19** |
| BD | BrainGENIE (10 PCs) | S-PrediXcan | 0.25 | 8.18 | 1.37E-16 | **3.70E-16** |
| BD | Blood | BrainGENIE (40 PCs) | -0.14 | -8.17 | 1.60E-16 | **4.17E-16** |
| BD | BrainGENIE (5 PCs) | S-PrediXcan | 0.25 | 8.15 | 1.87E-16 | **4.68E-16** |
| BD | BrainGENIE (20 PCs) | S-PrediXcan | 0.21 | 7.50 | 3.27E-14 | **7.30E-14** |
| BD | BrainGENIE (40 PCs) | S-PrediXcan | 0.19 | 6.91 | 2.46E-12 | **5.17E-12** |
| BD | BrainGENIE (10 PCs) | BrainGENIE (40 PCs) | 0.06 | 2.64 | 4.20E-03 | **5.81E-03** |
| BD | BrainGENIE (5 PCs) | BrainGENIE (40 PCs) | 0.06 | 2.62 | 4.36E-03 | **5.94E-03** |
| BD | BrainGENIE (10 PCs) | BrainGENIE (20 PCs) | 0.04 | 1.89 | 2.96E-02 | **3.68E-02** |
| BD | BrainGENIE (5 PCs) | BrainGENIE (20 PCs) | 0.04 | 1.88 | 2.98E-02 | **3.68E-02** |
| BD | Blood | S-PrediXcan | 0.05 | 1.74 | 4.11E-02 | **4.96E-02** |
| BD | BrainGENIE (20 PCs) | BrainGENIE (40 PCs) | 0.02 | 0.91 | 1.81E-01 | 1.96E-01 |
| BD | BrainGENIE (5 PCs) | BrainGENIE (10 PCs) | 0.00 | 0.02 | 4.93E-01 | 4.98E-01 |
| SCZ | Blood | BrainGENIE (5 PCs) | -0.47 | -25.19 | 2.38E-140 | **2.50E-138** |
| SCZ | Blood | BrainGENIE (20 PCs) | -0.33 | -21.66 | 2.32E-104 | **8.11E-103** |
| SCZ | Blood | BrainGENIE (10 PCs) | -0.36 | -18.74 | 1.19E-78 | **2.50E-77** |
| SCZ | BrainGENIE (5 PCs) | S-PrediXcan | 0.51 | 18.16 | 5.62E-74 | **6.56E-73** |
| SCZ | Blood | BrainGENIE (40 PCs) | -0.26 | -17.16 | 2.70E-66 | **2.83E-65** |
| SCZ | BrainGENIE (20 PCs) | S-PrediXcan | 0.37 | 13.77 | 1.89E-43 | **1.32E-42** |
| SCZ | BrainGENIE (10 PCs) | S-PrediXcan | 0.40 | 13.72 | 3.80E-43 | **2.49E-42** |
| SCZ | BrainGENIE (40 PCs) | S-PrediXcan | 0.30 | 11.17 | 2.94E-29 | **1.29E-28** |
| SCZ | BrainGENIE (5 PCs) | BrainGENIE (40 PCs) | 0.20 | 11.09 | 7.21E-29 | **3.03E-28** |
| SCZ | BrainGENIE (5 PCs) | BrainGENIE (20 PCs) | 0.14 | 7.84 | 2.28E-15 | **5.56E-15** |
| SCZ | BrainGENIE (5 PCs) | BrainGENIE (10 PCs) | 0.11 | 5.39 | 3.49E-08 | **6.43E-08** |
| SCZ | BrainGENIE (10 PCs) | BrainGENIE (40 PCs) | 0.10 | 4.94 | 3.95E-07 | **7.02E-07** |
| SCZ | BrainGENIE (20 PCs) | BrainGENIE (40 PCs) | 0.06 | 4.01 | 3.06E-05 | **4.79E-05** |
| SCZ | BrainGENIE (10 PCs) | BrainGENIE (20 PCs) | 0.03 | 1.66 | 4.87E-02 | 5.75E-02 |
| SCZ | Blood | S-PrediXcan | 0.04 | 1.38 | 8.41E-02 | 9.49E-02 |

| Concordance with differential gene expression signals from PsychENCODE (RNA-seq) | | | | | | |
|---|---|---|---|---|---|---|
| **Disorder** | **Method 1** | **Method 2** | **Δ Pearson's *r*** | ***z*-value** | ***p*-value** | **FDRp** |
| ASD | Blood | BrainGENIE (20 PCs) | -0.31 | -22.88 | 3.47E-116 | **1.82E-114** |
| ASD | Blood | BrainGENIE (10 PCs) | -0.31 | -18.62 | 1.12E-77 | **1.68E-76** |
| ASD | Blood | BrainGENIE (40 PCs) | -0.25 | -18.44 | 3.08E-76 | **4.04E-75** |
| ASD | Blood | BrainGENIE (5 PCs) | -0.23 | -13.42 | 2.20E-41 | **1.29E-40** |
| ASD | BrainGENIE (20 PCs) | S-PrediXcan | 0.26 | 13.19 | 5.16E-40 | **2.85E-39** |
| ASD | BrainGENIE (10 PCs) | S-PrediXcan | 0.27 | 12.03 | 1.17E-33 | **6.16E-33** |
| ASD | BrainGENIE (40 PCs) | S-PrediXcan | 0.20 | 10.17 | 1.33E-24 | **4.52E-24** |
| ASD | BrainGENIE (5 PCs) | S-PrediXcan | 0.18 | 8.16 | 1.63E-16 | **4.17E-16** |
| ASD | BrainGENIE (5 PCs) | BrainGENIE (20 PCs) | -0.07 | -4.29 | 8.92E-06 | **1.51E-05** |
| ASD | BrainGENIE (20 PCs) | BrainGENIE (40 PCs) | 0.06 | 4.09 | 2.17E-05 | **3.51E-05** |
| ASD | BrainGENIE (5 PCs) | BrainGENIE (10 PCs) | -0.08 | -4.08 | 2.21E-05 | **3.52E-05** |
| ASD | BrainGENIE (10 PCs) | BrainGENIE (40 PCs) | 0.06 | 3.76 | 8.63E-05 | **1.29E-04** |
| ASD | Blood | S-PrediXcan | -0.05 | -2.47 | 6.71E-03 | **8.70E-03** |
| ASD | BrainGENIE (5 PCs) | BrainGENIE (40 PCs) | -0.02 | -0.99 | 1.62E-01 | 1.77E-01 |
| ASD | BrainGENIE (10 PCs) | BrainGENIE (20 PCs) | 0.01 | 0.41 | 3.40E-01 | 3.50E-01 |
| BD | BrainGENIE (20 PCs) | BrainGENIE (40 PCs) | 0.11 | 7.64 | 1.09E-14 | **2.59E-14** |
| BD | BrainGENIE (20 PCs) | S-PrediXcan | 0.14 | 6.90 | 2.57E-12 | **5.29E-12** |

| | | | | | | |
|---|---|---|---|---|---|---|
| BD | BrainGENIE (5 PCs) | BrainGENIE (20 PCs) | -0.10 | -5.50 | 1.94E-08 | **3.64E-08** |
| BD | BrainGENIE (10 PCs) | S-PrediXcan | 0.09 | 4.11 | 1.96E-05 | **3.27E-05** |
| BD | Blood | BrainGENIE (20 PCs) | -0.06 | -3.86 | 5.77E-05 | **8.90E-05** |
| BD | BrainGENIE (10 PCs) | BrainGENIE (40 PCs) | 0.07 | 3.73 | 9.73E-05 | **1.44E-04** |
| BD | Blood | S-PrediXcan | 0.08 | 3.59 | 1.65E-04 | **2.40E-04** |
| BD | Blood | BrainGENIE (40 PCs) | 0.05 | 3.10 | 9.54E-04 | **1.35E-03** |
| BD | BrainGENIE (5 PCs) | BrainGENIE (10 PCs) | -0.05 | -2.61 | 4.49E-03 | **6.05E-03** |
| BD | BrainGENIE (10 PCs) | BrainGENIE (20 PCs) | -0.05 | -2.52 | 5.86E-03 | **7.79E-03** |
| BD | Blood | BrainGENIE (5 PCs) | 0.04 | 1.95 | 2.55E-02 | **3.23E-02** |
| BD | BrainGENIE (5 PCs) | S-PrediXcan | 0.04 | 1.67 | 4.78E-02 | 5.70E-02 |
| BD | BrainGENIE (40 PCs) | S-PrediXcan | 0.03 | 1.28 | 1.01E-01 | 1.13E-01 |
| BD | Blood | BrainGENIE (10 PCs) | -0.02 | -0.88 | 1.89E-01 | 2.03E-01 |
| BD | BrainGENIE (5 PCs) | BrainGENIE (40 PCs) | 0.01 | 0.68 | 2.50E-01 | 2.62E-01 |
| SCZ | Blood | BrainGENIE (5 PCs) | -0.24 | -13.71 | 4.74E-43 | **2.93E-42** |
| SCZ | BrainGENIE (5 PCs) | S-PrediXcan | 0.23 | 10.41 | 1.07E-25 | **3.89E-25** |
| SCZ | Blood | BrainGENIE (10 PCs) | -0.18 | -10.21 | 9.41E-25 | **3.29E-24** |
| SCZ | Blood | BrainGENIE (20 PCs) | -0.14 | -10.05 | 4.61E-24 | **1.51E-23** |
| SCZ | BrainGENIE (10 PCs) | S-PrediXcan | 0.17 | 7.63 | 1.20E-14 | **2.80E-14** |
| SCZ | BrainGENIE (5 PCs) | BrainGENIE (40 PCs) | 0.13 | 7.59 | 1.59E-14 | **3.63E-14** |
| SCZ | Blood | BrainGENIE (40 PCs) | -0.11 | -7.44 | 4.95E-14 | **1.08E-13** |
| SCZ | BrainGENIE (20 PCs) | S-PrediXcan | 0.13 | 6.75 | 7.14E-12 | **1.44E-11** |
| SCZ | BrainGENIE (5 PCs) | BrainGENIE (20 PCs) | 0.10 | 5.56 | 1.33E-08 | **2.54E-08** |
| SCZ | BrainGENIE (40 PCs) | S-PrediXcan | 0.10 | 4.91 | 4.66E-07 | **8.15E-07** |
| SCZ | BrainGENIE (10 PCs) | BrainGENIE (40 PCs) | 0.07 | 4.09 | 2.12E-05 | **3.48E-05** |
| SCZ | BrainGENIE (5 PCs) | BrainGENIE (10 PCs) | 0.06 | 3.08 | 1.03E-03 | **1.44E-03** |
| SCZ | BrainGENIE (20 PCs) | BrainGENIE (40 PCs) | 0.04 | 2.51 | 5.99E-03 | **7.86E-03** |
| SCZ | BrainGENIE (10 PCs) | BrainGENIE (20 PCs) | 0.04 | 2.04 | 2.07E-02 | **2.65E-02** |
| SCZ | Blood | S-PrediXcan | -0.01 | -0.38 | 3.50E-01 | 3.57E-01 |
| **Concordance with differential gene expression signals from CommonMind Consortium (RNA-seq)** | | | | | | |
| **Disorder** | **Method 1** | **Method 2** | **Δ Pearson's *r*** | **z-value** | ***p*-value** | **FDRp** |
| SCZ | Blood | BrainGENIE (40 PCs) | -0.24 | -16.51 | 1.50E-61 | **1.43E-60** |
| SCZ | S-PrediXcan | BrainGENIE (40 PCs) | -0.37 | -16.36 | 1.93E-60 | **1.69E-59** |
| SCZ | S-PrediXcan | BrainGENIE (5 PCs) | -0.30 | -11.85 | 1.05E-32 | **4.99E-32** |
| SCZ | BrainGENIE (20 PCs) | BrainGENIE (40 PCs) | -0.16 | -11.27 | 8.95E-30 | **4.09E-29** |
| SCZ | S-PrediXcan | BrainGENIE (10 PCs) | -0.27 | -10.55 | 2.65E-26 | **9.94E-26** |
| SCZ | Blood | BrainGENIE (5 PCs) | -0.17 | -9.20 | 1.77E-20 | **5.32E-20** |
| SCZ | S-PrediXcan | BrainGENIE (20 PCs) | -0.21 | -8.84 | 4.79E-19 | **1.32E-18** |
| SCZ | Blood | BrainGENIE (10 PCs) | -0.14 | -7.40 | 7.01E-14 | **1.50E-13** |
| SCZ | Blood | S-PrediXcan | 0.14 | 5.87 | 2.16E-09 | **4.28E-09** |
| SCZ | BrainGENIE (10 PCs) | BrainGENIE (40 PCs) | -0.10 | -5.63 | 8.75E-09 | **1.70E-08** |
| SCZ | BrainGENIE (5 PCs) | BrainGENIE (20 PCs) | 0.10 | 5.26 | 7.21E-08 | **1.30E-07** |
| SCZ | Blood | BrainGENIE (20 PCs) | -0.07 | -4.78 | 8.71E-07 | **1.50E-06** |
| SCZ | BrainGENIE (5 PCs) | BrainGENIE (40 PCs) | -0.07 | -3.83 | 6.37E-05 | **9.69E-05** |
| SCZ | BrainGENIE (10 PCs) | BrainGENIE (20 PCs) | 0.06 | 3.49 | 2.44E-04 | **3.51E-04** |
| SCZ | BrainGENIE (5 PCs) | BrainGENIE (10 PCs) | 0.03 | 1.54 | 6.18E-02 | 7.21E-02 |

### Supplementary Figures

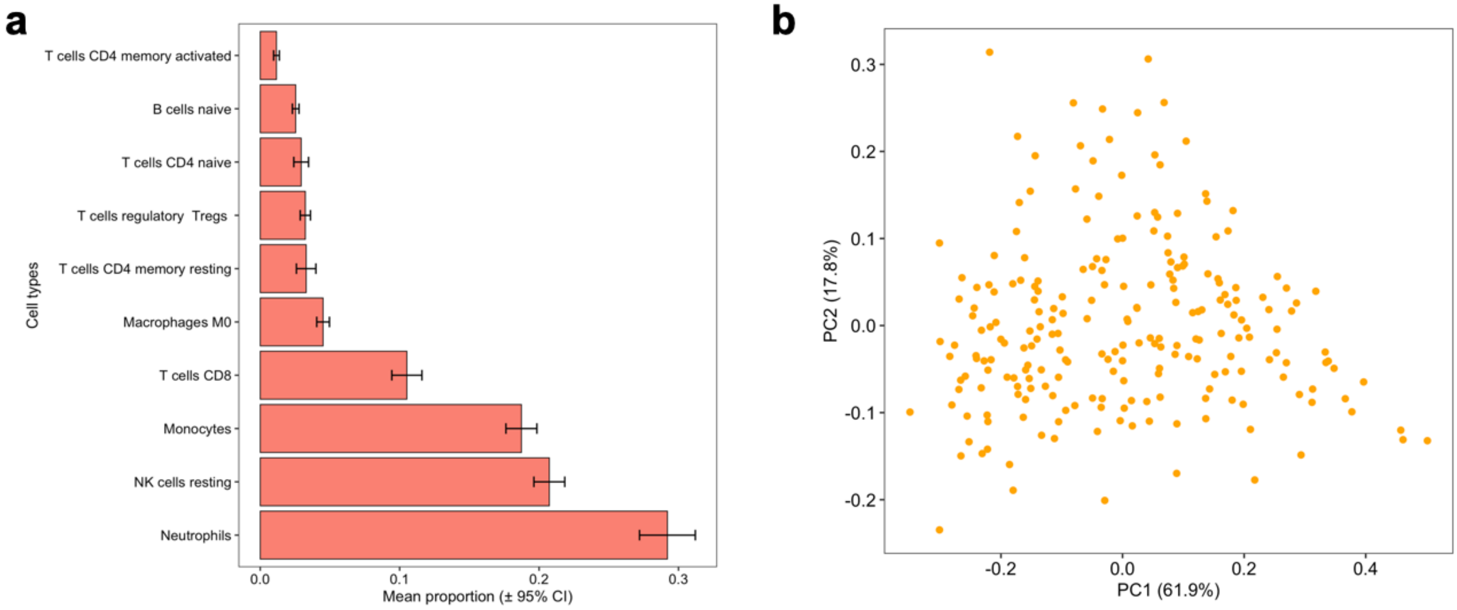

**Supplementary Figure 1.** Cell deconvolution analysis with *CIBERSORT* reveals the estimated abundance of circulating leukocytes in blood based on expression levels of cell-type marker genes assayed in whole blood from 225 donors in GTEx (v.8). (a) The estimated proportions of 10 circulating leukocytes are shown, which were averaged over 225 adult donors. The plot was limited to cell types with an average abundance ≥ 1%. (b) A scatterplot of donor scores for the first two components derived from principal components analysis (PCA) of estimated abundances of 22 leukocytes outputted from *CIBERSORT*. These component scores explained ~80% of the variance in leukocyte abundances between donors in GTEx. Donor scores for the top three principal components of blood cell abundances were regressed out of the whole blood gene expression data collected from the GTEx donors to adjust for unwanted variation in blood cell abundances that may have obfuscated (or spuriously created) cross-tissue gene expression correlations between blood and brain.

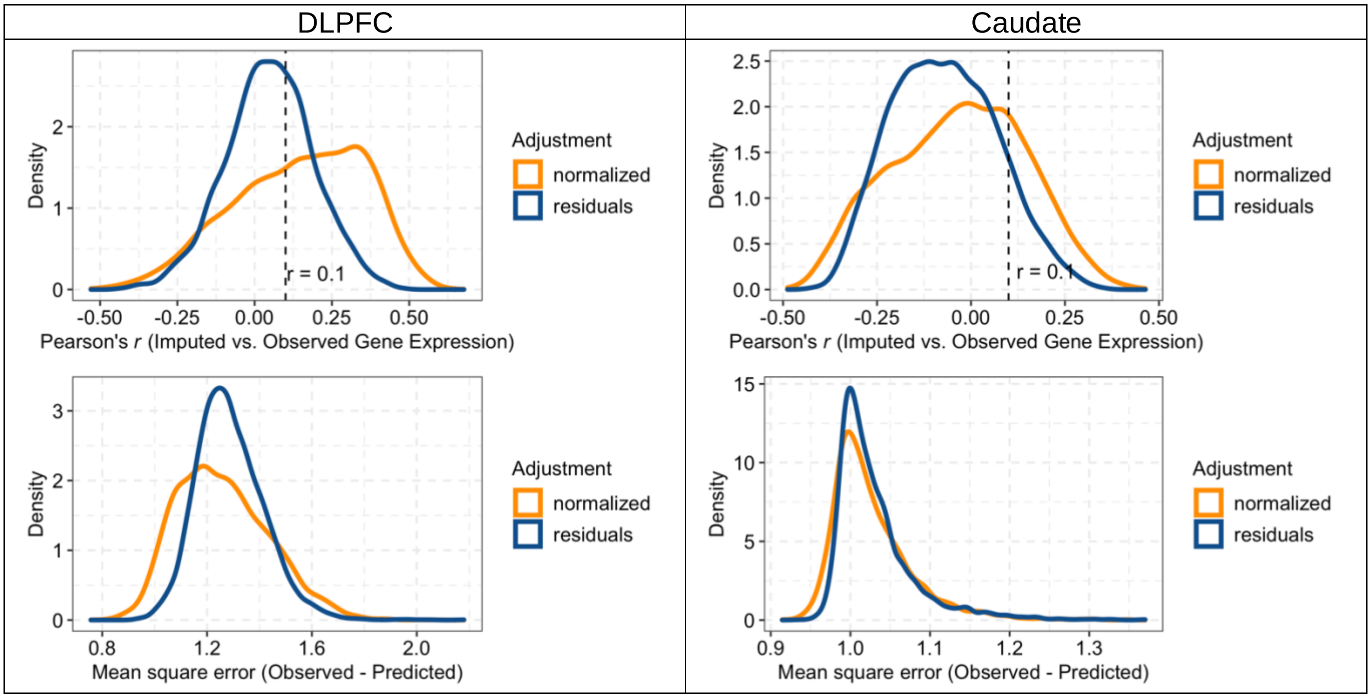

**Supplementary Figure 2**. Density plots of showing the impact of RNA-sequencing normalization on the accuracy (Pearson's *r*) and error (mean square error) of imputed brain gene expression levels generated by *BrainGENIE* in ROSMAP. We evaluated whether two normalization approaches to *postmortem* brain RNA-sequencing data derived from bulk brain tissue: 1) log-2 scaling of normalized gene expression values, and 2) log-2 scaling of normalized gene expression values *plus* derivation of residuals by partialling out variance explained by *postmortem* interval (PMI), RNA integrity number (RIN), library batch, and sequencing batch (DLPFC only). The "residuals" data were found to be over-corrected; *BrainGENIE*-imputed gene expression levels showed higher concordance (measured by Pearson's *r*) and lower error (measured by MSE) with the "normalized" data than the "residuals" data.

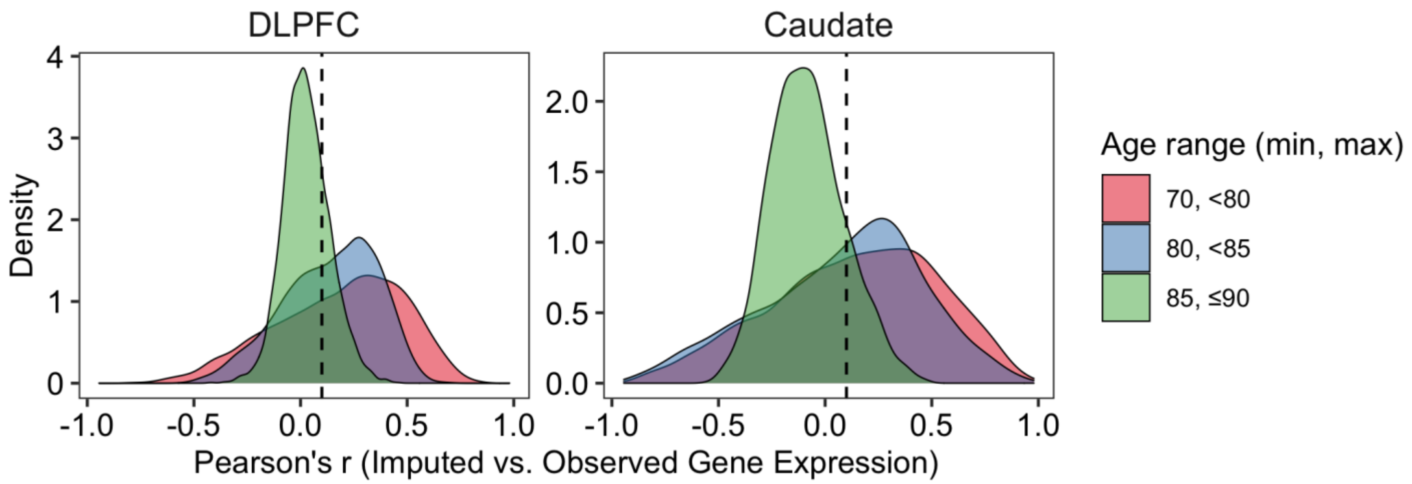

**Supplementary Figure 3**. A density plot showing the relationship between *BrainGENIE*'s imputation accuracy (measured by Pearson's *r*) and ages of individuals in ROSMAP at the time of blood draw for RNA extraction. The ages of individuals in ROSMAP were beyond the age range of the GTEx (v.8) samples used to derive imputation weights for *BrainGENIE*, which is suspected to play a role in diminishing accuracy in exceptional old adults. A vertical dotted line denotes Pearson's $r$=0.1. Individuals aged > 85 years old showed diminished imputation accuracy, thus were excluded from our *post-hoc* concordance analyses.

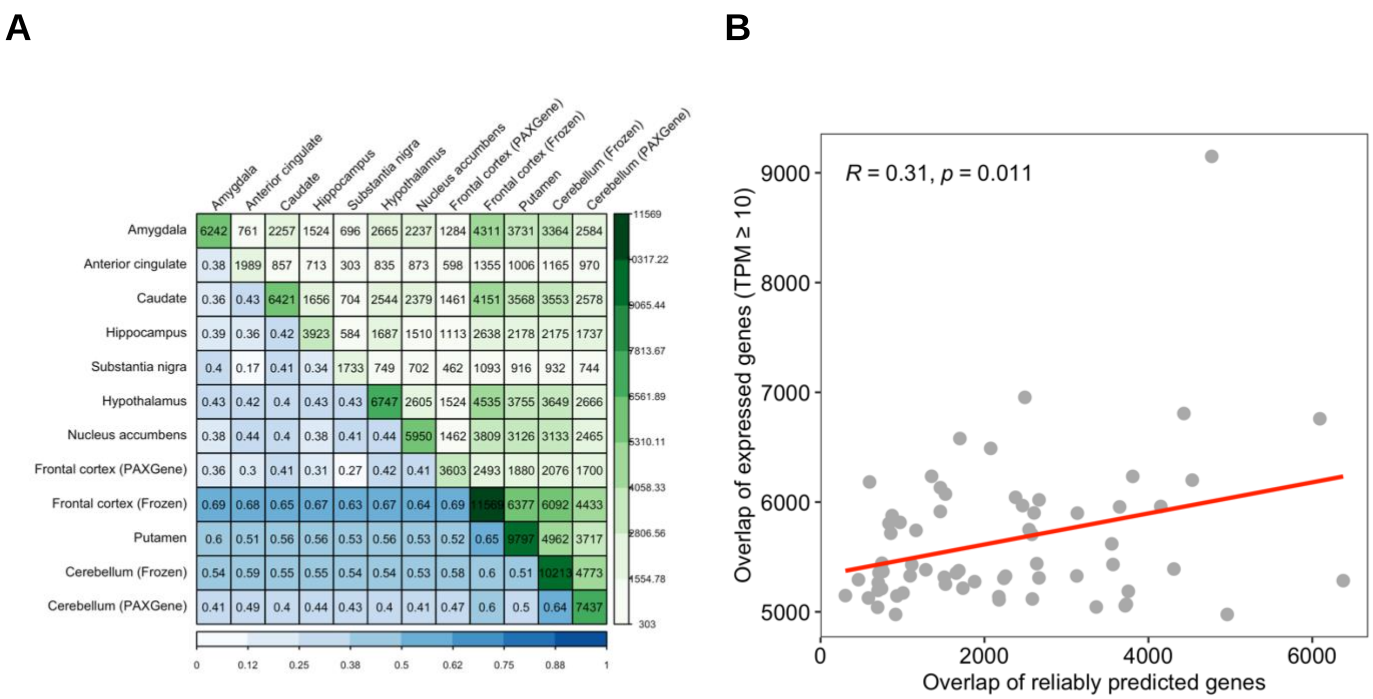

**Supplementary Figure 4**. *BrainGENIE* captures spatial relationships between brain tissues in GTEx. (**A**) A heatmap showing pairwise overlap of the number of genes for which *BrainGENIE* can reliably predict expression levels between 12 brain tissues in GTEx. The diagonal shows the number of genes reliably predicted in each of the 12 brain tissues. Values below the diagonal represent the proportion of genes from one brain tissue (the tissue with fewer reliably predicted genes) that overlapped in another brain tissue (the tissue with more reliably predicted genes). Values above the diagonal are the number of reliably predicted genes in common between pairs of brain tissues. (**B**) A scatter plot showing, on the vertical axis, the expected similarity between brain tissues in GTEx based on the number of overlapping genes expressed ≥ 10 transcripts per million (TPM). Plotted on the horizontal axis is the number of overlapping reliably predicted genes between brain tissues. The expected similarity between brain tissues was partially recapitulated by *BrainGENIE*.

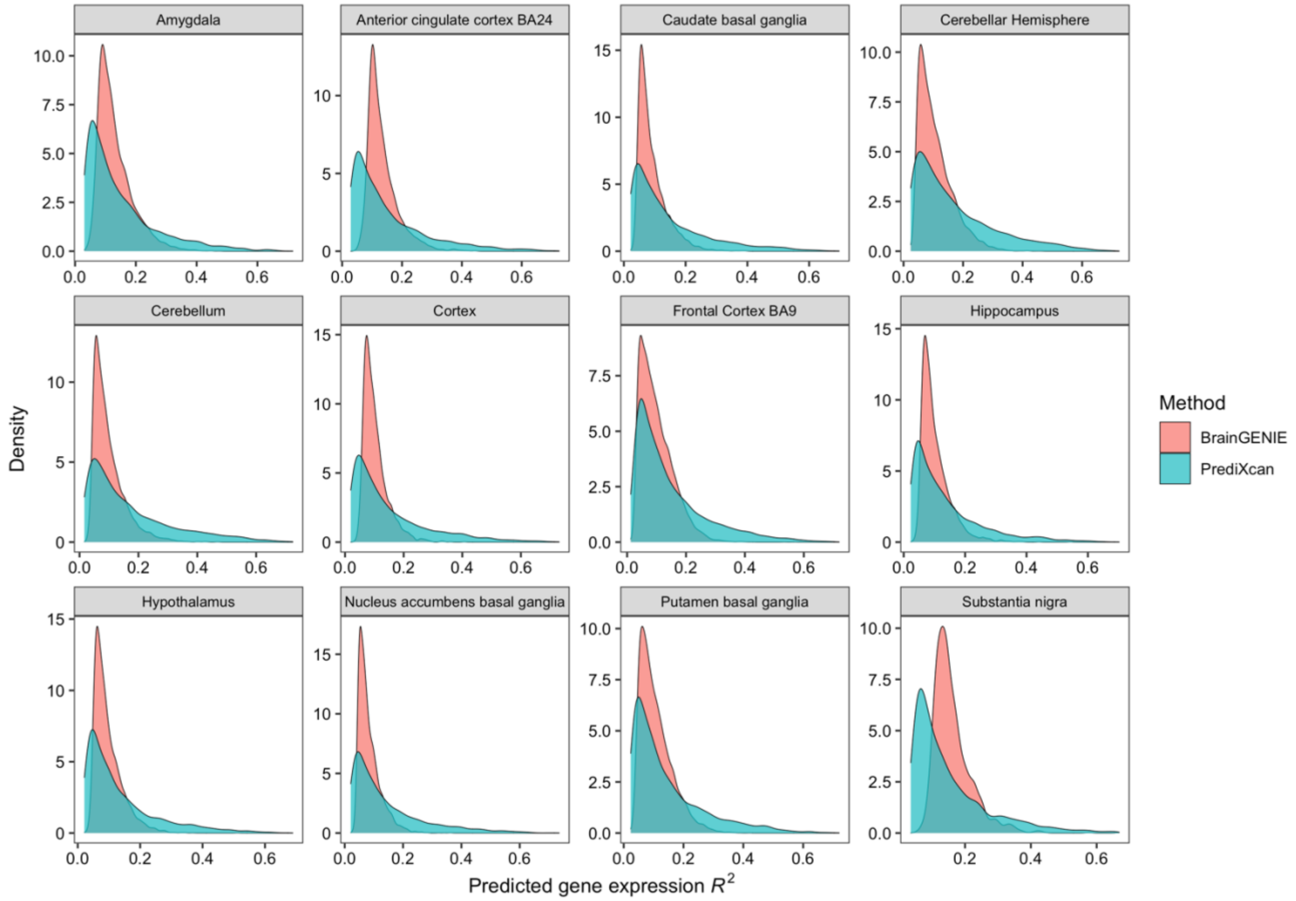

**Supplementary Figure 5**. Smoothed histograms showing the distribution of model prediction performances for all genes (measured by cross-validation $R^2$) whose expression levels in the brain were reliably predicted by *BrainGENIE* or *PrediXcan*.

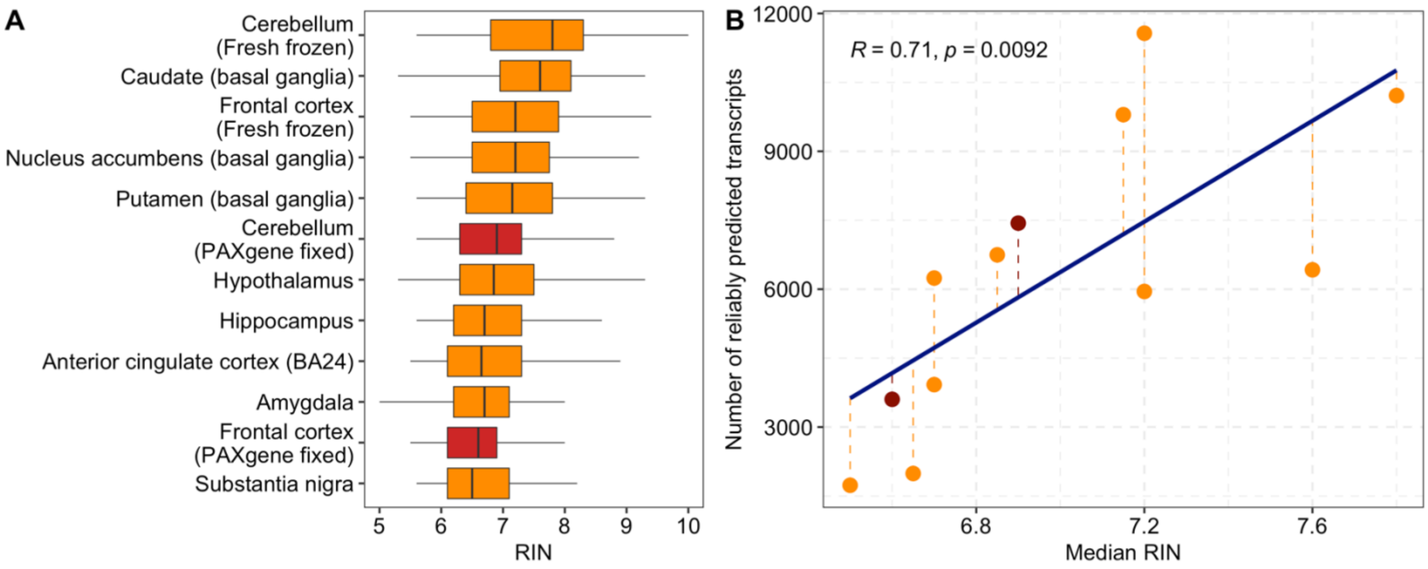

**Supplementary Figure 6**. Influence of RNA quality, measured by RNA integrity number (RIN), on the imputation performance of *BrainGENIE*. (**A**) The distribution of RIN is shown per brain tissue from GTEx (v8). Adjacently sampled brain tissues that were preserved in PAXgene tissue kits were highlighted in red; all other brain tissues were collected and fresh-frozen at the Univeristy of Miami Endowment Braink Bank. (**B**) The relationship between median RIN and number of genes with reliably predicted expression levels was plotted. Brain tissue samples preserved in PAXgene tissue kits were highlighted in red.

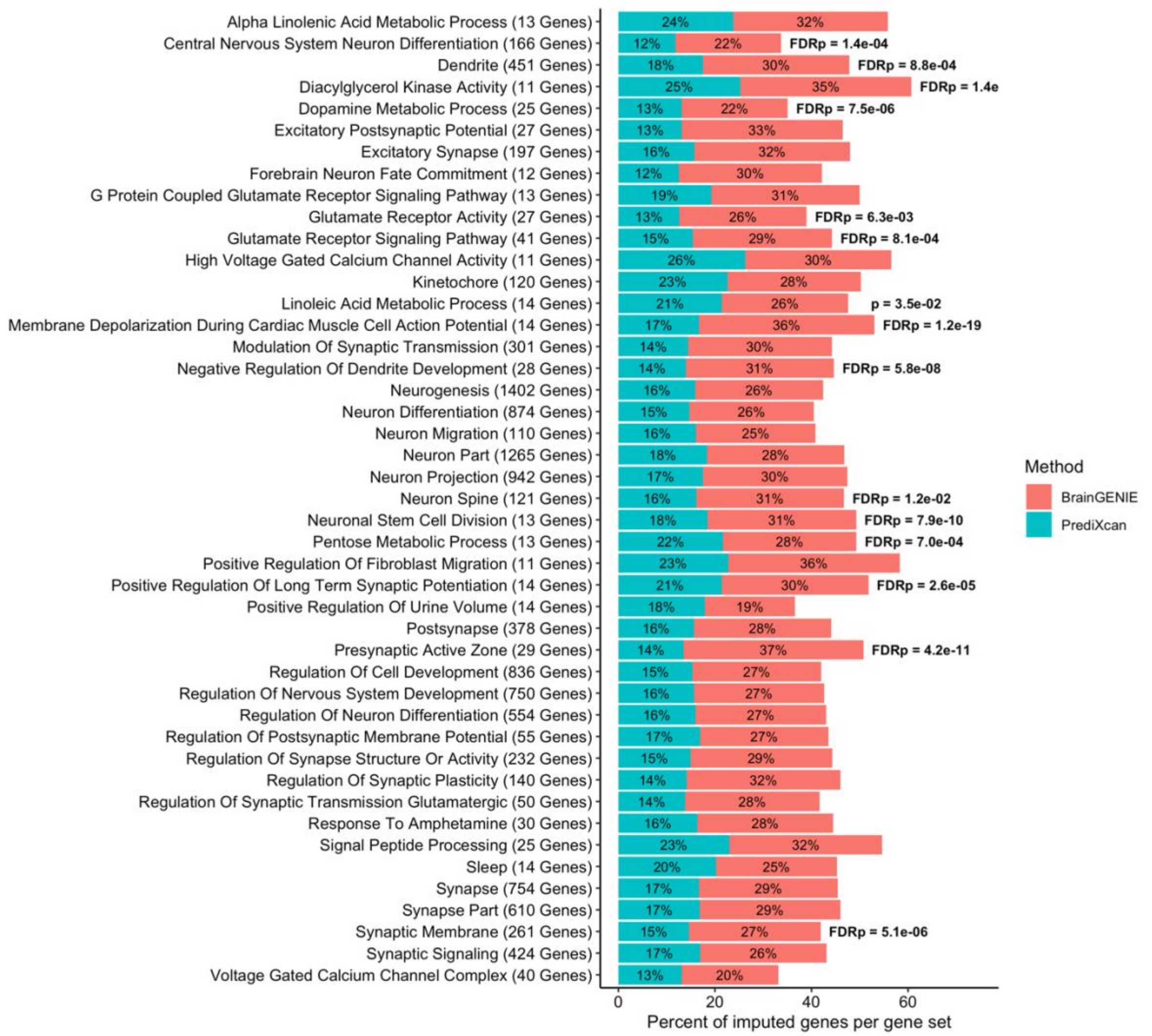

**Supplementary Figure 7**. A stacked bar plot showing the percent of genes reliably predicted by *BrainGENIE* or *PrediXcan* across 45 Gene Ontology (GO) gene sets known to be enriched with cross-disorder risk genes for eight neuropsychiatric disorders.[25] Bars were annotated with two-proportion Chi-square test *p*-values if *BrainGENIE* had imputed significantly more cross-disorder risk genes compared with *PrediXcan* (uncorrected *p*-value shown for tests yielding $p<0.05$, False-discovery-rated [FDR] adjusted *p*-value shown for tests reaching a FDRp<0.05).

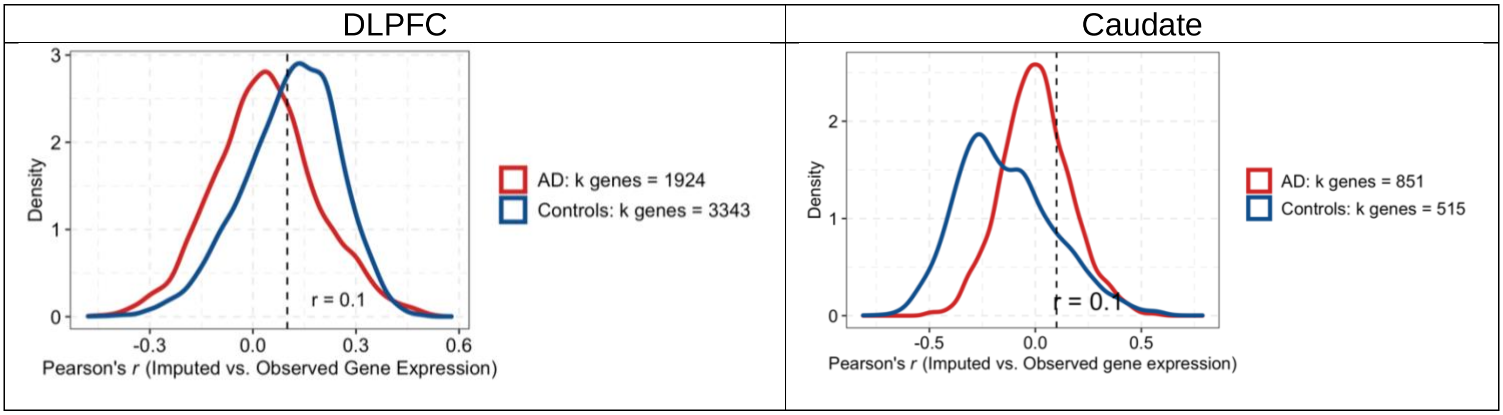

**Supplementary Figure 8**. Smoothed histograms showing the distribution of Pearson's *r* values measuring the accuracy of brain-imputed gene expression levels (produced by *BrainGENIE*) compared with observed *postmortem* DLPFC gene expression levels across subjects in the Religious Orders Study and Memory and Aging Project (ROSMAP). (**Left panel**) All individuals that contributed paired blood (i.e., *ex vivo* CD14+ monocytes) and *postmortem* DLPFC transcriptomes (*n*=109; AD *n*=54, Control *n*=55) were included. (**Right panel**) Individuals that contributed paired blood and *postmortem* caudate transcriptomes (*n*=50, AD=33, Control *n*=17) were included. The legends provide the number of genes that showed a minimum concordance of imputed *vs*. observed expression levels of Pearson's *r*≥0.1 per diagnostic group.

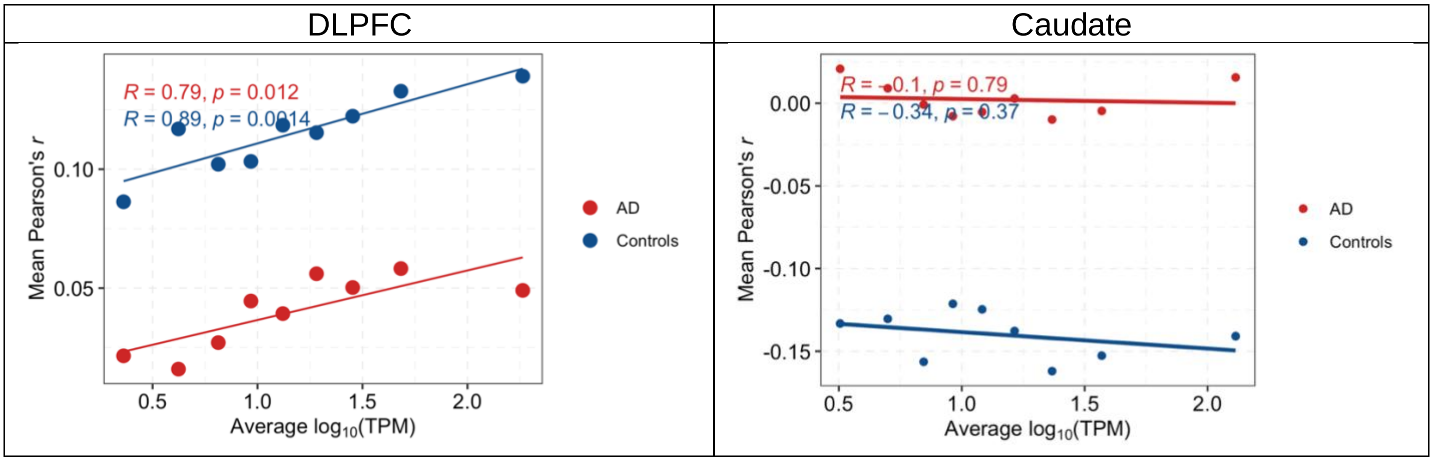

**Supplementary Figure 9**. Scatterplots showing the relationship of *BrainGENIE*'s imputation accuracy in ROSMAP (external test set) for the DLPFC transcriptome, measured by the square of Pearson's *r*, ($R^2$) relative to average expression levels of genes in $\log_{10}$-transformed transcripts per million (TPM) observed in brain tissue samples in our training dataset from GTEx (v.8). (**Left panel**) Average imputation accuracies for DLPFC were computed across 10 cut-points of gene expression containing approximately equal numbers of genes ($\log_{10}$TPM: 0.5, 1.8, 3.6, 5.7, 8.4, 12.1, 17.9, 27.2, 46.3, 175.5). The cut-points corresponded to the deciles of DLPFC $\log_{10}$TPM values. (**Right panel**) Average imputation accuracies for caudate were computed across 10 cut-points of gene expression containing approximately equal numbers of genes ($\log_{10}$TPM: 1.4, 3.2, 4.9, 6.7, 8.8, 11.5, 15.6, 22.7, 36.4, 119.5).

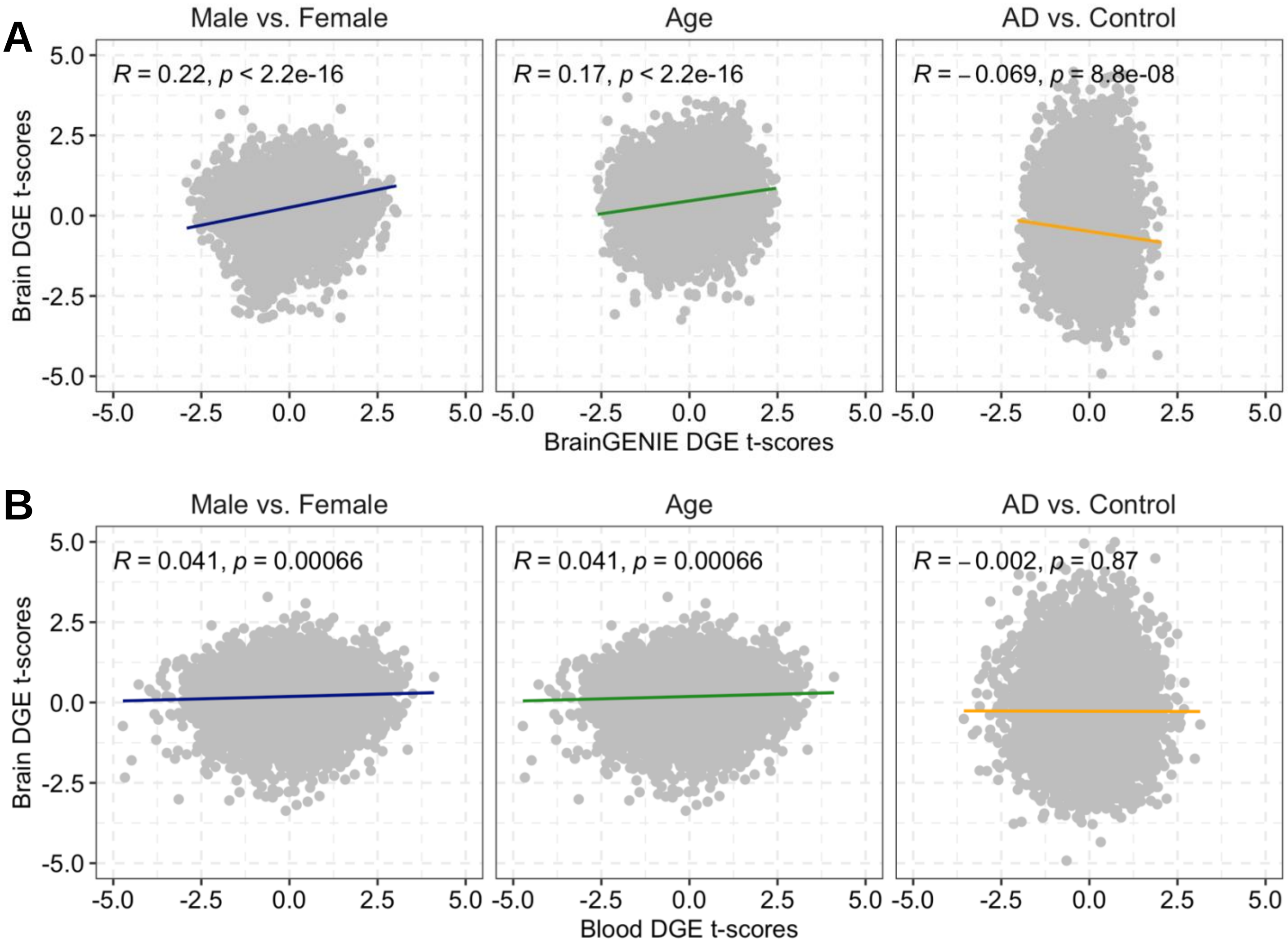

**Supplementary Figure 10**. Correlation analysis of differential gene expression (DGE) effect-sizes for age, sex, and Alzheimer's disease (AD) obtained from all ROSMAP with paired blood and DLPFC transcriptomes (*n*=109; AD *n*=54, Control *n*=55). Scatterplots show the paired samples correlation of differential gene expression effect-sizes to evaluate concordance of (**A**) frontal cortex-imputed and observed DLPFC transcriptomes, and (**B**) CD14+ purified monocyte and DLPFC transcriptomes.

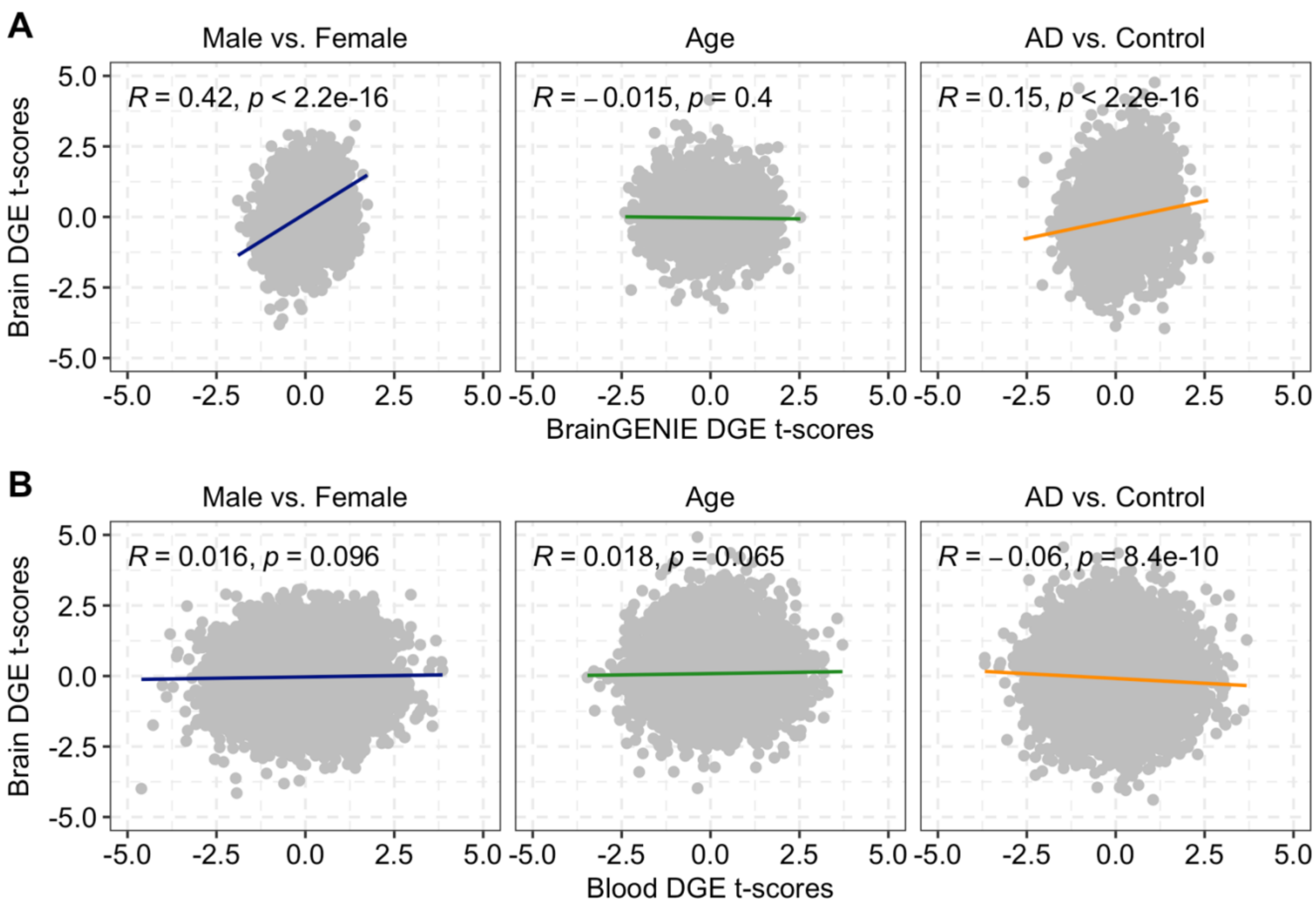

**Supplementary Figure 11**. Correlation analysis of differential gene expression (DGE) effect-sizes for age, sex, and Alzheimer's disease (AD) obtained from all ROSMAP with paired blood and caudate transcriptomes ($n$=50; AD $n$=33, Control $n$=17). Scatterplots show the paired samples correlation of differential gene expression effect-sizes to evaluate concordance of (**A**) caudate-imputed and observed caudate transcriptomes, and (**B**) CD14+ purified monocyte and caudate transcriptomes.
